# Upregulation of C_4_ characteristics does not consistently improve photosynthetic performance in intraspecific hybrids of a grass

**DOI:** 10.1101/2021.08.10.455822

**Authors:** Matheus E. Bianconi, Graciela Sotelo, Emma V. Curran, Vanja Milenkovic, Emanuela Samaritani, Luke T. Dunning, Lígia T. Bertolino, Colin P. Osborne, Pascal-Antoine Christin

## Abstract

C_4_ photosynthesis is thought to have evolved via intermediate stages, with changes towards the C_4_ phenotype gradually enhancing photosynthetic performance. This hypothesis is widely supported by modelling studies, but experimental tests are missing. Mixing of C_4_ components to generate artificial intermediates can be achieved via crossing, and the grass *Alloteropsis semialata* represents an outstanding system since it includes C_4_ and non-C_4_ populations. Here, we analyse F1 hybrids between C_3_ and C_4_, and C_3_+C_4_ and C_4_ genotypes to determine whether the acquisition of C_4_ characteristics increases photosynthetic performance. The hybrids have leaf anatomical characters and C_4_ gene expression profiles that are largely intermediate between those of their parents. Carbon isotope ratios are similarly intermediate, which suggests that a partial C_4_ cycle coexists with C_3_ carbon fixation in the hybrids. This partial C_4_ phenotype is associated with C_4_-like photosynthetic efficiency in C_3_+C_4_ x C_4_, but not in C_3_ x C_4_ hybrids, which are overall less efficient than both parents. Our results support the hypothesis that the photosynthetic gains from the upregulation of C_4_ characteristics depend on coordinated changes in anatomy and biochemistry. The order of acquisition of C_4_ components is thus constrained, with C_3_+C_4_ species providing an essential step for C_4_ evolution.

## Introduction

C_4_ photosynthesis is a complex adaptation that allows plants to sustain high photosynthetic rates in conditions that limit CO_2_ availability in the leaf, such as hot, dry and saline environments (Sage 2004). The C_4_ metabolism relies on a biochemical cycle operating within a specialized leaf anatomy, which concentrates CO_2_ around Rubisco, suppressing the energetically-costly photorespiratory pathway (Hatch 1987). The C_4_ trait is highly polyphyletic, but its multiple origins are mostly clustered in a few groups (Sage et al. 2011). The increased evolutionary accessibility to a C_4_ physiology in these groups has been associated with a number of ecological, anatomical and genetic features that acted as evolutionary enablers (Sage 2001; Christin et al. 2013; Moreno-Villena et al. 2018; Edwards 2019). The classic model of C_4_ evolution hypothesizes that, once these enablers are in place, an intermediate metabolism that relies on a photorespiratory CO_2_ pump is established (in ‘type I’ C_3_-C_4_ intermediates or ‘C2’ plants), and gradually complemented by a weak C_4_ cycle (in ‘type II’ C_3_-C_4_ *sensu* Edwards and Ku 1987, or ‘C3+C4’ plants *sensu* Dunning et al. 2017), which is subsequently optimized (Hylton et al. 1988; Monson and Moore 1989; Sage 2004). The transition through the intermediate stages has been predicted by mechanistic modelling studies, which support the idea that the sequential acquisition of components of the C_4_ trait successively increases fitness through increases in photosynthetic output (Heckmann et al. 2013, 2016; Mallmann et al. 2014). This model however currently lacks experimental support. Efforts to engineer the C_4_ trait into C_3_ crops have provided the first opportunities to study the effects of C_4_ components in isolation (Ishimaru et al. 1997; Taniguchi et al. 2008; Wang et al. 2017; Ermakova et al. 2021), but the recipient species do not necessarily possess the anatomical properties required for an efficient C_4_ metabolism. An alternative strategy is to segregate the components of the C_4_ trait via crosses between C_4_ and non-C_4_ plants (Simpson et al. In press).

Successful interspecific crosses between C_4_ and non-C_4_ plants have been reported particularly for the eudicot genera *Atriplex* (e.g. Björkman et al. 1969; Oakley et al. 2014) and *Flaveria* (e.g. Araus et al. 1990; Byrd et al. 1992; reviewed by Brown and Bouton 1993), with a recent interest in exploring this strategy to dissect the C_4_ trait (Oakley et al. 2014; Sultmanis 2018; Lin et al. 2021; Simpson et al. In press). Most of the diversity of C_4_ lineages however lies in monocots, and especially the grass family, with at least 22 independent origins of C_4_ photosynthesis (GPWG II 2012). The only existing crosses between photosynthetic types in monocots concerned C_3_ and C_3_-C_4_ parents (Bouton et al. 1986), and the grass *Alloteropsis semialata* offers a particularly suitable system to complement these studies, as it includes not only C_3_ and C_4_ (Ellis 1974a), but also C_3_+C_4_ individuals (Lundgren et al. 2016). While different photosynthetic types of *A. semialata* can hybridize (Bianconi et al. 2020), the effects of different crosses remain to be described.

*Alloteropsis semialata* has a paleotropical distribution, with C_3_ and C_3_+C_4_ populations in Southern and Central/Eastern regions of Africa, respectively, and C_4_ individuals occurring across Africa, South and Southeast Asia, and Oceania (Lundgren et al. 2015; Olofsson et al. 2016; Bianconi et al. 2020; Olofsson et al. 2021). The photosynthetic types in *A. semialata* are associated with distinct genetic lineages (Lundgren et al. 2015; Olofsson et al. 2016), which initially diverged around 3 Mya (Bianconi et al. 2020). Although the ranges of C_4_ and non-C_4_ lineages geographically overlap in some regions of Africa, with C_3_+C_4_ and C_4_ in Central/Eastern Africa, and C_3_ and C_4_ in Southern Africa, genetic analyses have shown that natural hybrids are rare (Olofsson et al. 2016; Bianconi et al. 2020; Olofsson et al. 2021). The lack of a clear hybrid zone has been partially explained by the differences in ploidy levels among photosynthetic types where they are found in sympatry (Olofsson et al. 2021). However, the broad ranges of C_4_ and C_3_+C_4_ diploids overlap, and they have been coexisting for at least one million years (Lundgren et al. 2015; Bianconi et al. 2020). Besides ploidy differences, the low frequency of natural hybridization might be associated with pre- and post-zygotic barriers, such as possible hybrid depression associated with pleiotropic costs of upregulating components of C_4_ photosynthesis without a fully functional C_4_ metabolism in place (Olofsson et al. 2021). Experimental crosses between C_4_ and non-C_4_ individuals of *A. semialata* provide an opportunity to test this hypothesis.

Here we generate hybrids between C_3_ and C_4_ (C_3_ x C_4_), and C_3_+C_4_ and C_4_ (C3+C_4_ x C_4_) accessions of *A. semialata* and analyse their phenotype to test the hypotheses that components of the C_4_ trait are additive, and that hybrids rank between their non-C_4_ and C_4_ parents in terms of photosynthetic performance. Our study confirms that crosses between photosynthetic types in *A. semialata* are viable in experimental conditions, and shows that the C_4_ metabolism is disrupted in the hybrids despite the significant upregulation of anatomical and biochemical components of the C_4_ trait.

## Material and Methods

### Plant material and growth conditions

Crosses between accessions of *Alloteropsis semialata* (R. Br.) Hitchc. (Poaceae) from distinct geographical origins were generated from parental plants collected in the wild as seeds or cuttings (Table S1) and grown in greenhouse conditions at the Arthur Willis Environment Centre, University of Sheffield (UK). Initial attempts of controlled cross-pollination using pollination bags to isolate hand-pollinated inflorescences were unsuccessful. For this reason, we adopted a non-controlled pollination strategy, in which plants were allowed to receive pollen from any other plant in the greenhouse, with subsequent determination of the pollen parent via genotyping (see below). The resulting seeds were collected while still attached to the mother plant, and subsequently germinated on Petri dishes before being potted. Here, a total of 31 seedlings from seven mother plants (two C_3_, two C_3_+C_4_ and three C_4_) were obtained. Subsequent genotyping of the seedlings identified multiple F1 hybrids between parents with distinct photosynthetic types, including 14 C_3_ x C_4_ and six C_3_+C_4_ x C_4_ individuals (Table S1). The remaining seedlings consisted of 10 C_4_ x C_4_ crosses, and an additional C_3_+C_4_ plant resulting from self-pollination (Table S2). Five C_3_ x C_4_ individuals grew to maturity and were sampled for DNA analyses but died before being phenotyped, and another two died before being sampled for RNA sequencing and leaf gas-exchange analyses (Table S1). For the phenotypic analyses, we included the mother plants, the known and possible pollen parents (i.e. similar C_4_ accessions that could not be distinguished by the genotyping analyses; see Table S1), and seven additional accessions of *A. semialata* (three C_3_, two C_3_+C_4_ and two C_4_) that were growing alongside them were added to increase the phenotypic diversity within photosynthetic types (Table S1). All plants were grown in 11-L, free-draining pots containing a 2:1 mix of M3 compost (Levington) and perlite (Sinclair) under well-watered conditions, and were fertilized once every three months with a NK fertilizer 16-0-5 containing Iron (Evergreen Extreme Green Lawn Food). Plants were grown on a 12-hour day/night cycle, with metal halide lamps providing supplementary light (additional 200 μmol m^-2^ s^-1^ at bench level), 25/20°C day/night temperature, ambient CO_2_ concentration and relative humidity between 30% and 60%.

### Genotyping

We used three genetic markers to determine the genetic lineage of the pollen parent of each plant. These markers correspond to genomic regions (∼ 500 bp long) with sufficient variation to distinguish between different photosynthetic types and geographical groups, and were selected from those previously assembled to determine the origin of allopolyploids in *A. semialata* (Bianconi et al. 2020). Custom primers were designed to PCR amplify the target genes (Table S3). Genomic DNA (gDNA) of the putative crosses was isolated from fresh leaves using the DNeasy Plant Mini Kit (Qiagen, Hilden, Germany). PCR reactions contained ca. 10-40 ng of gDNA template, 5 µl of 5X GoTaq Flexi reaction buffer (Promega, Madison, WI, USA), 2 mM of MgCl2, 0.08 mM of dNTPs, 0.2 µM of each primer and 0.5 unit of GoTaq polymerase (Promega) in a total volume of 25 µl. The PCR mixtures were initially incubated in a thermocycler for 2 min at 94°C followed by 35 cycles consisting of 30 s at 94°C (denaturation), 1 min at 48°C (annealing) and 1 min at 72°C (elongation). Amplicons were cleaned using Exo-SAP-IT (Affymetrix, Santa Clara, CA, USA), and Sanger-sequenced at the Core Genomic Facility at the University of Sheffield. Sequencing chromatograms were individually inspected for heterozygous sites that are polymorphic among genetic groups corresponding to distinct photosynthetic types, since the presence of such heterozygous sites indicates a F1 hybrid between photosynthetic types. Because the maternal origin was known in advance (the maternal plant was the one on which the seed was collected), the information from the alternative bases was used to narrow down the pollen parent to a genetic lineage (Table S2). The hybrid origin of the plants was subsequently confirmed using RNAseq data (see below).

### Stable carbon isotopes

Carbon isotope composition was determined from the central portion of fully expanded leaf blades. Samples were dried in silica gel, and subsequently ground to a fine powder using a TissueLyzer II (Qiagen). Carbon isotope analysis was conducted on 1-2 mg of leaf sample using an ANCA GSL preparation module coupled to a Sercon 20-20 stable isotope ratio mass spectrometer (PDZ Europa, Cheshire, UK). Carbon isotopic ratios (δ^13^C, in ‰) were reported relative to the standard Pee Dee Belemnite (PDB). Values of δ^13^C higher than -16‰ indicate that the plants grew using C_4_ photosynthesis (O’Leary 1988; Stata et al. 2019). δ^13^C values of additional accessions of *A. semialata* that were previously phenotyped and identified as C_3_, C_3_+C_4_ or C_4_ (Lundgren et al. 2016, 2019; Dunning et al. 2017, 2019a) were retrieved from Lundgren et al. (2015, 2016).

### Leaf anatomy

Leaf cross sections were obtained from the central portion of fully expanded leaf blades. Fresh samples were initially dehydrated in an ethanol series from 70% to 100% EtOH, and resin-infiltrated with Technovit 7100 (Heraeus Kulzer GmbH, Wehrheim, Germany) following the manufacturer’s instructions. Cross sections 7-10 μm thick were obtained using a microtome (Leica RM 2245, Leica Biosystems Nussloch GmbH, Nussloch, Germany), and stained with Toluidine Blue O (Sigma-Aldrich, St. Louis, MO, USA). An Olympus BX51 microscope coupled to an Olympus DP71 camera (Olympus Corporation, Tokyo, Japan) was used to photograph the cross sections. Leaf anatomical traits were measured using ImageJ v.1.51q (Schneider et al. 2012) in one cross section per individual. Measurements were taken following Lundgren et al. (2019) within segments, where the segment was defined as the area between two consecutive secondary veins (i.e. veins with large metaxylem vessels; see Fig. S1). Segments that were adjacent to the mid vein or to the lateral margins of the cross section were avoided to maintain consistency between samples. The variables measured here included: leaf thickness (at the leftmost secondary vein in the segment), the number of minor veins (4^th^ and 5^th^ order veins), vein density, distance between edges of consecutive bundle sheaths (BSD), width of inner (IS) and outer sheath (OS), and the areas of epidermis (including bulliform cells, Epd), mesophyll (M), extraxylary fibres (Fb), bundle sheath (separately for IS and OS) and vascular tissue (V; see Fig. S1 for an illustrated key for each variable). Areas were expressed relative to the total area of the segment before analysis, except IS and OS, which were expressed relative to the mesophyll area (IS/M and OS/M). A principal component analysis (PCA) was performed on these 12 leaf anatomical variables using the function *prcomp* in R v. 3.6.3 (R Core Team 2020).

### Leaf transcriptome

Leaf mRNA was isolated and sequenced as previously described (Dunning et al. 2019a). In short, the distal halves of fully expanded leaves were sampled in the middle of the light period, flash-frozen in liquid N2 and kept at -80°C. All samples were collected on the same day. Total RNA was extracted using the RNeasy Plant Mini Kit (Qiagen, Germany) with an on-column DNA digestion step (RNase-Free DNase Set; Qiagen). A total of 20 RNA-Seq libraries (one per individual) were prepared with the TruSeq RNA Library Preparation Kit v2 (Illumina, San Diego, CA, USA) using 0.5 μg of starting RNA and aiming at a median insert size of ∼ 155 bp (standard fragmentation protocol). Libraries were paired-end sequenced (read length = 100 bp) on 1/24 of a single lane of an Illumina HiSeq 2500 flow cell in rapid mode (with four additional samples from an unrelated project) at the Sheffield Diagnostic Genetics Service. Raw sequence reads were filtered to remove adaptor contamination and low-quality reads (i.e. < 80% of bases with Phred score > 20) using NGSQCToolkit v. 2.3.3 (Patel and Jain 2012). Reads were further trimmed from the 3’ end to remove bases with Phred score < 20. The quality of the filtered data was assessed using FastQC v. 0.11.9 (Andrews 2010).

Transcript abundance was quantified by mapping the filtered reads to a reference dataset consisting of coding sequences from multiple transcriptomes of *A. semialata* retrieved from Dunning et al. (2017) and modified by Bianconi et al. (2018). In short, this dataset consists of 5,540 groups of homologous genes that are common to the Panicoideae grasses (the subfamily that includes *Alloteropsis*), each group containing all paralogs detected in the transcriptomes of *A. semialata* (total of 12,234 groups of co-orthologs). We used this dataset as it includes manually curated sets of co-orthologs of 23 gene families known to have a function in the C_4_ biochemistry (Bianconi et al. 2018). This curated gene set increases the read mapping accuracy when paralogs exist with a high sequence similarity. Furthermore, this dataset includes the sequences of paralogs of core C_4_ genes that are absent from the reference genome of *A. semialata*, such as four different genes for the enzyme phosphoenolpyruvate carboxylase (PEPC) known to be highly expressed in other *A. semialata* accessions (Dunning et al. 2017, 2019a). Filtered reads were mapped to the reference dataset using Bowtie2 v. 2.3.5 (Langmead and Salzberg 2012) with default parameters. Read counts are reported here in reads per million mapped reads (RPM). To visualize the diversity among samples, a PCA on the global patterns of gene expression was conducted using the function *prcomp* in R after log2-transforming the RPM values. Only genes with transcript abundance > 10 RPM in at least five samples were used for the PCA. To test whether the global patterns of gene expression were consistent when the whole gene set of *A. semialata* was considered, we repeated the analysis using the chromosome-level genome assembly of this species (Dunning et al. 2019b) as reference for read mapping. Finally, to verify whether the expression level differences between photosynthetic types were consistent with previous reports for *A. semialata*, we quantified transcript abundance (using the co-orthologous gene set as reference) for 20 additional accessions (four C_3_, six C_3_+C_4_ and 10 C_4_) retrieved from two previous RNAseq studies (Dunning et al. 2017, 2019a).

To explore the global differences in gene expression between hybrids and the parental types, we performed a differential expression analysis on the dataset that was generated using the complete genome of *A. semialata* as reference for read mapping. Here, due to insufficient number of replicates for C_3_ and C_3_+C_4_ plants, we restricted our analyses to the comparisons between the two hybrid types, and between each of these and the C_4_ group. We then investigated the main metabolic functions associated with the set of differentially expressed genes from each comparison (see full description of the global RNAseq analyses on the Supplementary Methods).

Finally, to confirm the paternity of the putative hybrids, we genotyped the plants using the RNAseq dataset mapped to the reference genome of *A. semialata*. Here we also included individuals of *A. semialata* from the RNAseq studies of Dunning et al. (2017, 2019a) to increase the genetic diversity of the dataset. Read alignment files were filtered to remove duplicates using Picard Tools v. 1.102 (https://broadinstitute.github.io/picard/), and variants were called on all individuals combined using BCFtools v. 1.9 (Li et al. 2009). Variant sites were filtered using VCFtools v. 0.1.17 (Li et al. 2009) to remove low quality genotype calls (i.e. quality score < 30 and read depth < 3), indels, and sites with more than 10% missing data, which resulted in an initial set of 521,929 single nucleotide polymorphisms (SNPs). BCFtools was then used to retain only multiallelic sites in which all hybrids were heterozygous and all potential parents were homozygous, which resulted in 15,406 SNPs that were then used for the genotyping tests. First, to determine the type of cross (i.e. whether C_3_ x C_4_ or C_3_+C_4_ x C_4_), we identified SNPs that were common to all individuals of the same photosynthetic type and exclusive to them. Sites with more than 25% missing data within each photosynthetic group were removed, and a total of 114 SNPs were retained (Table S4). We then tested, for each SNP, whether hybrid individuals had one C_4_ allele and one allele that was either C_3_ or C_3_+C4, according to the type of cross (Table S4). With the confirmation of the type of cross, we then narrowed down the genetic lineage of the pollen parent (as the mother is already known). Only one individual representing each genetic lineage was retained for this analysis (Table S5). First, we used the 15,406 SNP set to identify all singletons of each potential pollen parent, which in this case were defined as homozygous genotypes that were unique to a single individual among all individuals with the same photosynthetic type. A total of 7,779 singletons were identified (minimum of 287 per potential pollen parent; Table S5). We then verified, for each singleton, which hybrid individuals carried that same allele, and counted the number of positive matches (Table S5). All genotyping analyses were conducted using R (scripts are provided as Supplementary Files).

### Leaf gas-exchanges

The photosynthetic response to intercellular CO_2_ concentration (*A*/*Ci* curve) was measured using two portable gas-exchange systems Li-6400XT (Li-Cor, Lincoln, NE, USA). *A*/*Ci* curves were measured within the first six hours of the photoperiod on the widest fully expanded leaf at a block temperature (*T_block_*) of 25 °C, flow rate = 300 μmol s^-1^, photosynthetic photon flux density (*PPFD*) = 1500 μmol m^-2^ s^-1^, and were started after both net CO_2_ uptake (*A*) and stomatal conductance (*gS*) reached steady-state at reference CO_2_ = 400 μmol mol^-1^. Reference CO_2_ was then changed in a stepwise manner to 250, 150, 120, 100, 85, 70, 50, 35, 400, 600, 800, 1000 and 1200 μmol mol^-1^.

Readings were automatically logged after 2 to 3 minutes of leaf acclimation to each CO_2_ level. The CO_2_ compensation point (CCP) and maximum carboxylation efficiency (CE) were calculated following Bellasio et al. (2016). In this approach, the CO_2_-dependence of *A* is described by an empirical non-rectangular hyperbola, and allows for the estimation of parameters irrespective of the photosynthetic physiology, which is therefore a suitable approach for hybrids (Bellasio et al. 2016). Water-use efficiency (WUE) was calculated as the ratio between *A* and *gS* at steady-state at reference CO_2_ = 400 μmol mol^-1^.

Stomatal density on the abaxial side of the leaves was quantified using leaf impressions. Dental resin (ImpressPlus Wash Light Body, Perfection Plus Ltd., Totton, UK) was applied to the central portion of fully expanded leaves, and clear nail varnish was applied to the set resin impression. Images were captured from the nail varnish impressions at 20 X using an Olympus BX51 microscope coupled to an Olympus DP71 camera, and stomatal density was quantified on 0.38 mm² fields using ImageJ.

## Results

### Carbon isotopic ratios are intermediate between the parents

Carbon isotopic ratios (δ^13^C) of all hybrids between C_4_ and non-C_4_ accessions were below the range of C_4_ values in *A. semialata* (Fig. 1; Table S1). However, δ^13^C was consistently higher in the hybrids than in their non-C_4_ parents, with average increases of 2.8 ‰ in C_3_ x C_4_ and 5.4 ‰ in C_3_+C_4_ x C_4_ hybrids relative to their C_3_ and C_3_+C_4_ parents, respectively. The δ^13^C ranged between -27.7 and - 25.3 ‰ in C_3_ x C_4_, and -21.4 and -18.5 ‰ in C_3_+C_4_ x C_4_ hybrids.

**Fig. 1.**
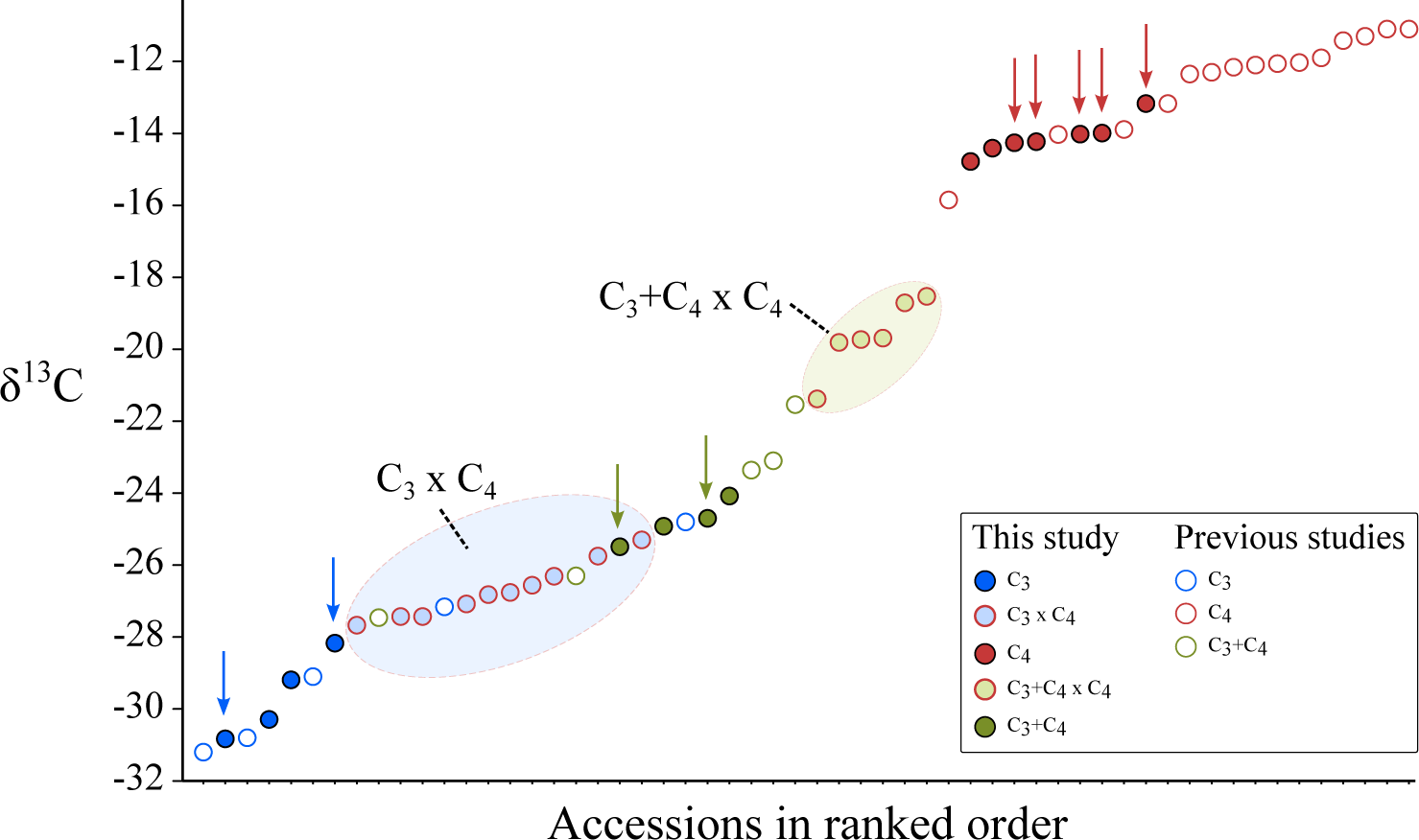
Distribution of carbon isotopic ratios (δ^13^C) in *Alloteropsis semialata*. δ^13^C values of accessions of *A. semialata* that are not part of this study were retrieved from Bianconi et al. (2020). Blue, green and red arrows indicate δ^13^C values of C_3_, C_3_+C_4_ and C_4_ known or potential parents of the hybrids analysed in this study.

### Leaf anatomy is additive in the hybrids

C_4_ and non-C_4_ leaves in *A. semialata* are mainly distinguished by the presence of minor veins in C_4_ accessions (Lundgren et al. 2019). Here, minor veins were observed in all hybrids, except two C_3_ x C_4_ individuals (Table S6). The presence of chloroplasts in the inner sheath of all hybrids is suggested by the strong and consistent staining patterns in these cells, which are similar to the patterns observed in C_4_ and C_3_+C_4_ *A. semialata*, but not in C_3_ accessions, where the staining is very weak (Fig. 2A). The first component of a PCA on 12 quantitative anatomical traits explains 57.1% of the total variation and separates the accessions per photosynthetic types (Fig. 2B). C_4_ individuals are associated with negative values of the first component, which correspond to increased vein density (including minor veins) and increased abundance of bundle sheath tissue (IS and OS) relative to the mesophyll (Fig. 2B). C_3_+C_4_ and C_3_ accessions partially overlap in the PCA space, and are associated with positive values of the first component. Hybrid individuals are intermediate between their respective parents along this first component. The second principal component explains 13.9% of the variance, and is correlated with leaf thickness (r = 0.88, *p* < 0.001), width of inner bundle sheath cells (ISW; r = 0.64, *p* < 0.001) and epidermis area (Epd; r = -0.54, *p* < 0.001), but these variables do not differentiate photosynthetic or cross types. The proportion of inner bundle sheath relative to mesophyll (IS/M) was on average doubled in C_3_ x C_4_ hybrids compared to C_3_ accessions, and was within the range of C_3_+C4, but the values were still 62% lower than the C_4_ average (Fig. 2C). The distance between consecutive bundle sheaths (BSD) in C_3_ x C_4_ and C_3_+C_4_ x C_4_ hybrids was on average 49% and 55% smaller than in C_3_ and C_3_+C_4_ plants, respectively (Fig. 2D). The mean width of IS cells was similar between C_3_+C_4_ and C_4_ accessions, and values were on average 48% higher than in C_3_, but there differences among photosynthetic types or between hybrids and the respective parental types were not significant (Fig. 2E). OS cells were however significantly smaller in C_4_ than in C_3_ accessions, with hybrids having intermediate values that were not significantly different from those of C_3_+C_4_ accessions (Fig. 2F). In the hybrids, the variation in BSD was largely explained by vein density (R^2^ = 0.92, *p* < 0.001), with no significant effects of OS or IS widths. The IS/M ratio, on the other hand, was not significantly associated with vein density, but with both OS and IS width (R^2^ = 0.75, *p* < 0.001). Overall, C_3_ x C_4_ hybrids were not significantly different from C_3_+C_4_ in relation to the quantitative anatomical traits analysed here, except for BSD, which was significantly reduced in C_3_ x C_4_ as a result of the presence of some minor veins in most accessions. In C_3_+C_4_ x C_4_ hybrids, a quantitatively distinct phenotype was generated, with trait values ranking between C_3_+C_4_ and C_4_ plants. Overall, our results show that leaf anatomical traits in *A. semialata* are mostly inherited in an additive manner.

**Fig. 2.**
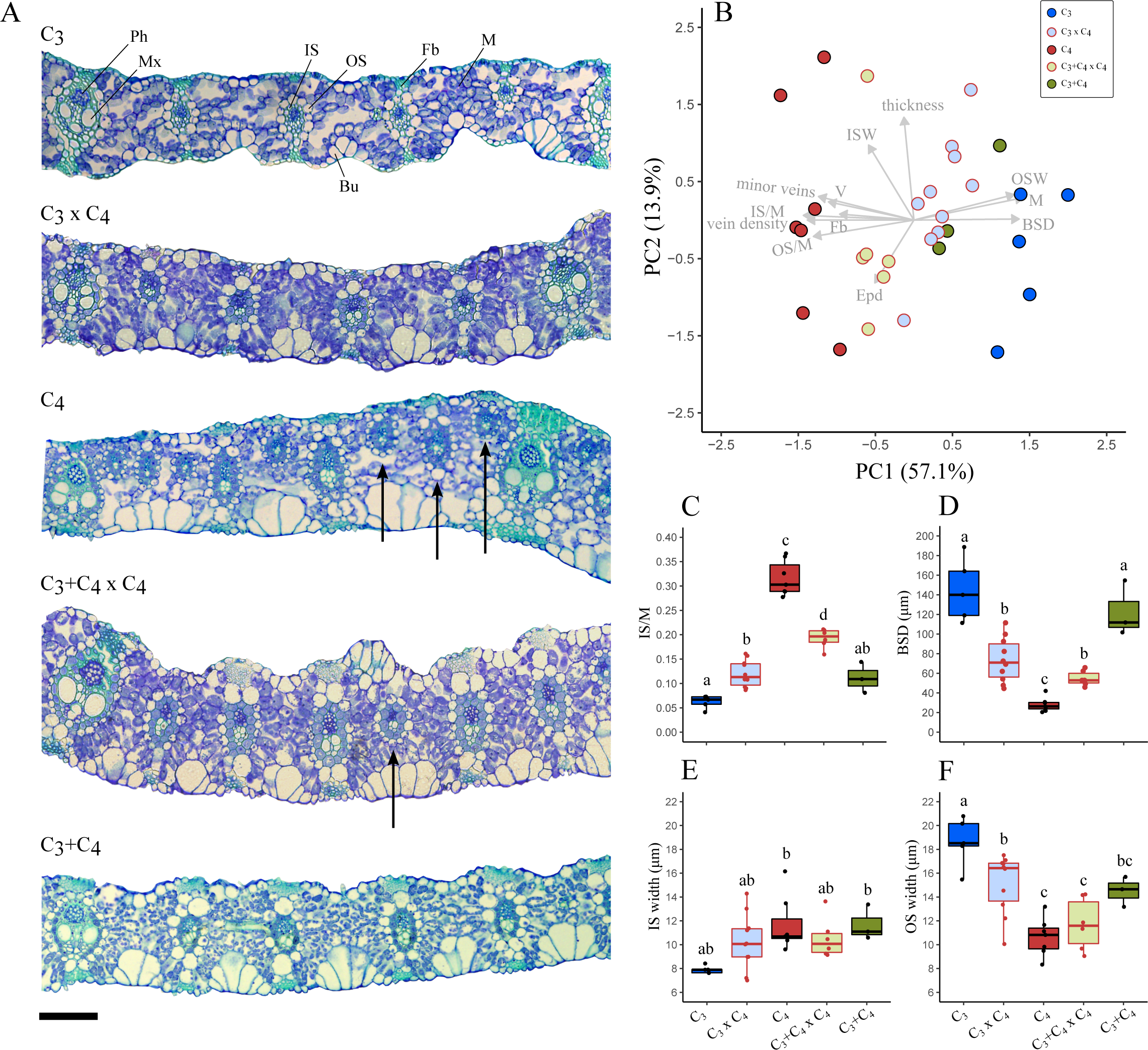
Leaf anatomy of F1 hybrids and the parental photosynthetic types in *Alloteropsis semialata* (A) Representative cross sections of C_3_ (RSA9), C_3_ x C_4_ (H08), C_4_ (TPE1-10), C_3_+C_4_ x C_4_ (H11) and C_3_+C_4_ (TAN1602-03c) accessions. Arrows indicate minor veins (i.e. fourth and fifth order veins). Scale bar = 100 µm. (B) Principal component analysis of selected leaf anatomical variables (C) Proportion of inner sheath to mesophyll area (IS/M). (D) Distance between consecutive bundle sheaths (BSD). (E) Mean inner sheath cell width. (F) Mean outer sheath cell width. For (C-F), different lower-case letters indicate statistical differences between groups (ANOVA, *p* < 0.05 post-hoc Tukey HSD; n ≥ 4). Bu = bulliform cells, BSD = distance between consecutive bundle sheaths, Epd = proportion of epidermis area, Fb = extraxylary fibres (= sclerenchyma girder), IS = inner bundle sheath (= mestome sheath), ISW = IS width, M = mesophyll, Mx = metaxylem, OS = outer bundle sheath, OSW = OS width, Ph = phloem, V = proportion of vascular tissue area.

### Gene expression

Global gene expression patterns were assessed via a PCA on 7,482 genes (Fig. 3A). The first principal component mostly separated three sister C_3_+C_4_ x C_4_ plants (H06, H11, and H23) from the other individuals, and accounted for 24.7% of the variation. Photosynthetic types were clearly separated by the second principal component, which accounted for 16.8% of the variation and placed the hybrids as intermediate to the photosynthetic types of their parents. Similar clustering patterns were observed when transcript abundance was quantified for the complete gene set extracted from the genome of *A. semialata* (Fig. S2).

**Fig. 3.**
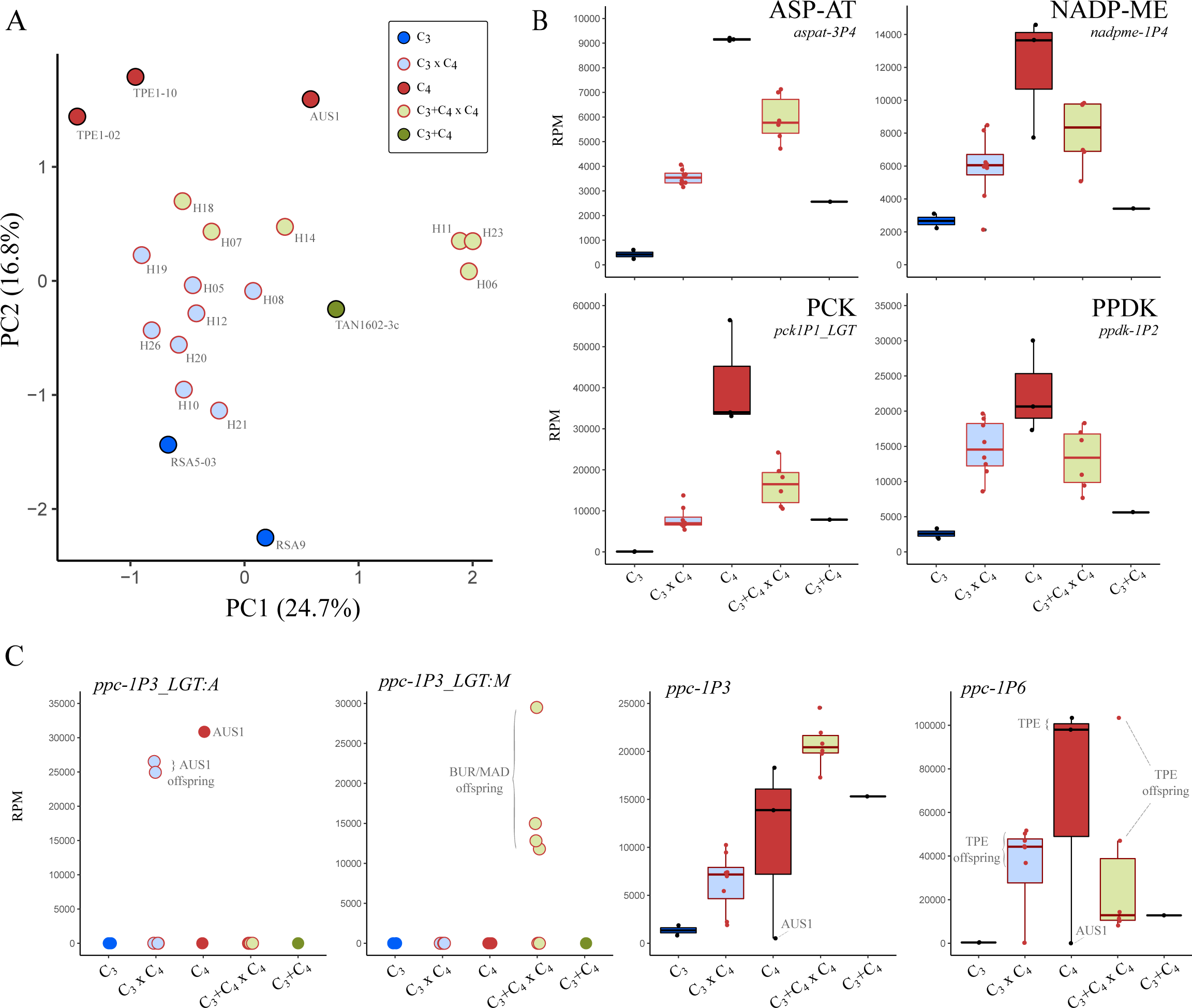
Leaf gene expression of F1 hybrids and the parental photosynthetic types in *Alloteropsis semialata*. (A) Principal component analysis on 7,482 genes. (B) Transcript abundance in reads per million mapped reads (RPM) of selected genes encoding core C_4_ enzymes: aspartate aminotransferase (ASP-AT, gene *aspat-3P4*), NADP-dependent malic enzyme (NADP-ME, gene *nadpme-1P4*), phosphoenolpyruvate carboxykinase (PCK, gene *pck1P1_LGT*), pyruvate phosphate dikinase (PPDK, gene *ppdk-1P2*). (C) Transcript abundance of selected phosphoenolpyruvate carboxylase (PEPC) genes.

Transcript abundance of five genes encoding key enzymes for the C_4_ biochemistry, and which are known to be upregulated in C_4_ plants of *A. semialata* (Dunning et al. 2019a), were analysed in detail (Fig. 3B,C; Table S7). The patterns of transcript abundance of C_3_, C_3_+C_4_ and C_4_ plants are highly similar to those reported by Dunning et al. (2017, 2019a), where a larger sample size was used (Fig. S3). Transcript abundance of the genes encoding the C_4_-specific forms of the enzymes aspartate aminotransferase (ASP-AT, gene *aspat-3P4*), NADP-malic enzyme (NADP-ME, gene *nadpme-1P4*) and pyruvate phosphate dikinase (PPDK, gene *ppdk-1P2*) were four- to 17-fold higher in C_4_ than in C_3_ plants, with the transcript abundances of the C_3_+C_4_ plant slightly above those of C_3_ plants (Fig. 3B). In each case, the transcript abundance of the hybrids was intermediate to the abundance of the photosynthetic types of the parents, showing that these expression patterns are mostly additive. C_3_ accessions of *A. semialata* lack the gene copy encoding phosphoenolpyruvate carboxykinase (PCK) that is highly expressed in C_4_ and C_3_+C_4_ accessions (*pck-1P1_LGT*; Olofsson et al. 2016); the transcript abundance of this gene reaches 20% of the mean C_4_ values in C_3_ x C_4_ hybrids, and 40% in C_3_+C_4_ x C_4_.

Out of the nine genes encoding the key C_4_ enzyme phosphoenolpyruvate carboxylase (PEPC) that have been identified in *A. semialata* accessions (Dunning et al. 2017), only five were reported to be highly expressed in leaves of some C_4_ accessions (Christin et al. 2012; Dunning et al. 2017, 2019a). One of these genes, *ppc-1P3_LGT:A* is only present in the C_4_ lineage from Oceania, which is represented here by an Australian accession (AUS1) that is the pollen parent of two C_3_ x C_4_ hybrids (H19 and H21; Table S5); both plants have expression levels of this gene that are similar to the Australian parent (Fig. 3C). Only one of the other two laterally-acquired PEPC copies is present in the plants analysed here (*ppc-1P3_LGT:M*), and it is highly expressed (> 10,000 RPM) in four C_3_+C_4_ x C_4_ individuals that are offspring from the C_4_ accessions from Burkina Faso or Madagascar (H06, H11, H14 and H23; Table S5; Fig. 3C), which have been previously shown to be part of the only C_4_ lineage to carry this gene (Olofsson et al. 2016). The expression levels of *ppc-1P3* are similar across C_3_+C_4_ and C_4_ accessions, and are on average ten-fold higher than in C_3_ accessions (except the C_4_ AUS1, which has a non-functional copy of this gene; Olofsson et al. 2016); in the C_3_+C_4_ x C_4_ hybrids, however, the transcript abundance of this gene is on average 42% higher than the median of C_3_+C_4_ and C_4_ accessions (Fig. 3C). Finally, the gene *ppc-1P6* reaches high expression levels in C_4_ accessions (except in AUS1, where it is absent; Dunning et al. 2019b), but is disproportionally upregulated in the C_4_ Taiwanese lineage (TPE; Dunning et al. 2017), where it reaches levels that are up to 100-fold higher than in C_3_ plants, and at least twice higher than the values in other C_4_ and C_3_+C_4_ accessions. Here, the *ppc-1P6* copy reaches high levels in the C_3_ x C_4_ and C_3_+C_4_ x C_4_ offspring of TPE, mostly ranging from 10,000 to 60,000 RPM, except for one C_3_+C_4_ x C_4_ which has a transcript abundance similar to TPE (∼ 100,000 RPM).

We examined transcript abundance of selected protein-coding genes with a role in the photorespiratory metabolism. We observed reduced transcript levels of the genes encoding the peroxisomal enzymes flavin mononucleotide-dependent glycolate oxidase (GOX, gene *glo-1P2*) and glutamate:glyoxylate aminotransferase (GGT, gene *ggat-1P6*) in C_3_+C_4_ x C_4_ hybrids relative to C_3_ and C_3_+C_4_ accessions, with average values that are similar to or below those of C_4_ accessions (Fig. S4). The genes encoding the proteins P-, T-, L- and H- that compose the mitochondrial multienzyme system glycine decarboxylase (GDC, genes *gldp1P*, *gcvt-1P1*, *lpd-1P2*, and *gdh-2P2*, respectively) had on average 63% lower transcript levels in C_4_ than in C_3_ accessions; in C_3_ x C_4_ and C_3_+C_4_ x C_4_ hybrids, transcript levels of these four genes were on average 12% and 33% lower than in C_3_, respectively, suggesting additivity (Fig. S4; Table S8). Multiple genes encoding a serine hydroxymethyltransferase (SHMT) are found in the genome of *A. semialata* (genes *shm-1P1*, *shm-2P2*, *shm-3P*3 and *shm-3P4*; Dunning et al. 2019a), but none had patterns of transcript abundance similar to those observed for GDC; in fact, one of the genes (*shm-3P*3) had higher average transcript abundance in C_4_ than in C_3_ accessions. Finally, the chloroplast genes glycerate 3-kinase (GLYK, gene *glyk-1P1*) and 2-phosphoglycolate (2-PG) phosphatase (PGLP, gene *pglp-2P2*) had overlapping transcript abundance ranges across hybrid and photosynthetic types. Finally, a gene encoding a GOLDEN2-LIKE transcription factor that has been previously demonstrated to be involved in the redistribution of organelles among cell types (Wang et al. 2017) is expressed at higher levels in C_4_ than in C_3_ and C_3_+C_4_ accessions (Fig. S4L; Table S8). This gene is at intermediate levels in the C_3_ x C_4_ hybrids, but largely variable among C_3_+C_4_ x C_4_ individuals.

We identified genes differentially expressed (DE) between hybrid types, and between these and C_4_ accessions (Table S9). We found a total of 4,104 genes differentially expressed between C_3_ x C_4_ and C_3_+C_4_ x C_4_ hybrids (adjusted *p*-value < 0.05), out of which 21 were related to C_4_ photosynthesis and four to photorespiration (Table S9). This complete DE gene set is reduced to 206 when only higher count genes (base mean > 500) and with larger differences (log2fold change > 1) are considered (Table S9). Among this reduced gene set, we found seven C_4_-related genes, but none related to photorespiration (Fig. S5; Table S9). The individual comparisons between C_3_ x C_4_ or C_3_+C_4_ x C_4_ and C_4_ accessions identified 1,925 and 2,727 significant DE genes, respectively. These numbers are reduced to 110 and 181 when the same filters described above are applied (Table S9). In this reduced set, we found 13 C_4_-related genes that were differentially expressed between hybrids and C_4_ accessions, three of these common to both C_3_ x C_4_ vs C_4_, and C_3_+C_4_ x C_4_ vs C_4_ comparisons, namely PCK (*pck-1P1_LGT*), NADP-malate dehydrogenase (NADP-MDH, gene *nadpmdh-1P2*), and soluble inorganic pyrophosphatase (PPA, gene *ppa-2P1*). While *nadpmdh-1P2* was downregulated in the C_4_ group, the two others had increased transcript abundance in C_4_ relative to both hybrid types (Fig. S5). Finally, we found four genes related to photorespiration in the DE gene set between C_3_ x C_4_ and C_4_ accessions, three of which were upregulated in C_3_ x C_4_, namely the genes encoding the GDC proteins T- (*gcvt-1P1*), P- (*gldp1P*) and L- (*lpd-1P2;* Fig. S5). The remaining gene, which encode the enzymes SHMT (*shm-3P*3) was downregulated in both C_3_ x C_4_ and C_3_+C_4_ x C_4_ hybrids relative to C_4_ accessions. Gene ontology (GO) analyses did not detect any significantly enriched cellular processes for any of the full DE gene sets. However, the reduced DE gene set from the comparison between hybrid types was significantly enriched with genes related to the photosystems I and II, oxidoreductase activity and 1,3-beta-D-glucan synthesis (Table S10). In the reduced gene set from comparisons between each of the hybrid types and the C_4_ type, GO analyses indicated a significant enrichment for genes associated with C_4_ photosynthesis (particularly malate metabolism), transmembrane transport, and photosystems I and II (Table S10).

Overall, the analyses of expression profiles show that the inheritance of expression levels of C_4_ genes and those involved in the photorespiratory pathway is largely additive. In almost all cases, the transcript abundance of the hybrids ranks between those of the parental lineages.

### Evidence of reduced photosynthetic performance in the hybrids

Photosynthetic performance was assessed from CO_2_ response curves (A/Ci) and steady-state measurements at ambient CO_2_ (Fig. S6; Tables S11 and S12). Photosynthetic rates in the hybrids ranked between the parents at low *Ci* levels, but the differences between hybrids and parents were progressively reduced as CO_2_ levels increased, until they were equal to their non-C_4_ parents at *Ci* of 100 in C_3_ x C_4_, and 250 µmol mol^-1^ in C_3_+C_4_ x C_4_ hybrids (Fig. 4). The C_3_ and C_3_+C_4_ accessions had on average similar values of maximum carboxylation efficiency (CE), and these were 57% lower than in C_4_ accessions (Fig. 5A; Table S11). Despite the larger variation among individuals, C_3_+C_4_ x C_4_ hybrids had CE values that were on average 83% and 96% higher than in C_3_ and C_3_+C_4_ accessions, respectively, and these were not significantly different from the average values of C_4_ accessions. In C_3_ x C_4_ hybrids, however, CE was consistently lower than in C_3_ accessions, and reached the lowest mean values of all photosynthetic types/crosses. The CO_2_ compensation point (CCP) ranged between 43 and 48 µmol mol^-1^ in C_3_ accessions, and between 5 and 15 µmol mol^-1^ in C_3_+C_4_ individuals (Fig. 5B). All C_4_ had CCP below 10 µmol mol^-1^. In the hybrids, CCP was below 13 µmol mol^-1^ in C_3_+C_4_ x C_4_ individuals, and ranged between 6 and 40 µmol mol^-1^ in C_3_ x C_4_ hybrids. CCP estimates varied substantially among A/*Ci* curves in C_3_ x C_4_ individuals (Fig. 5B), but mean values were 52% lower than in C_3_ plants.

**Fig. 4.**
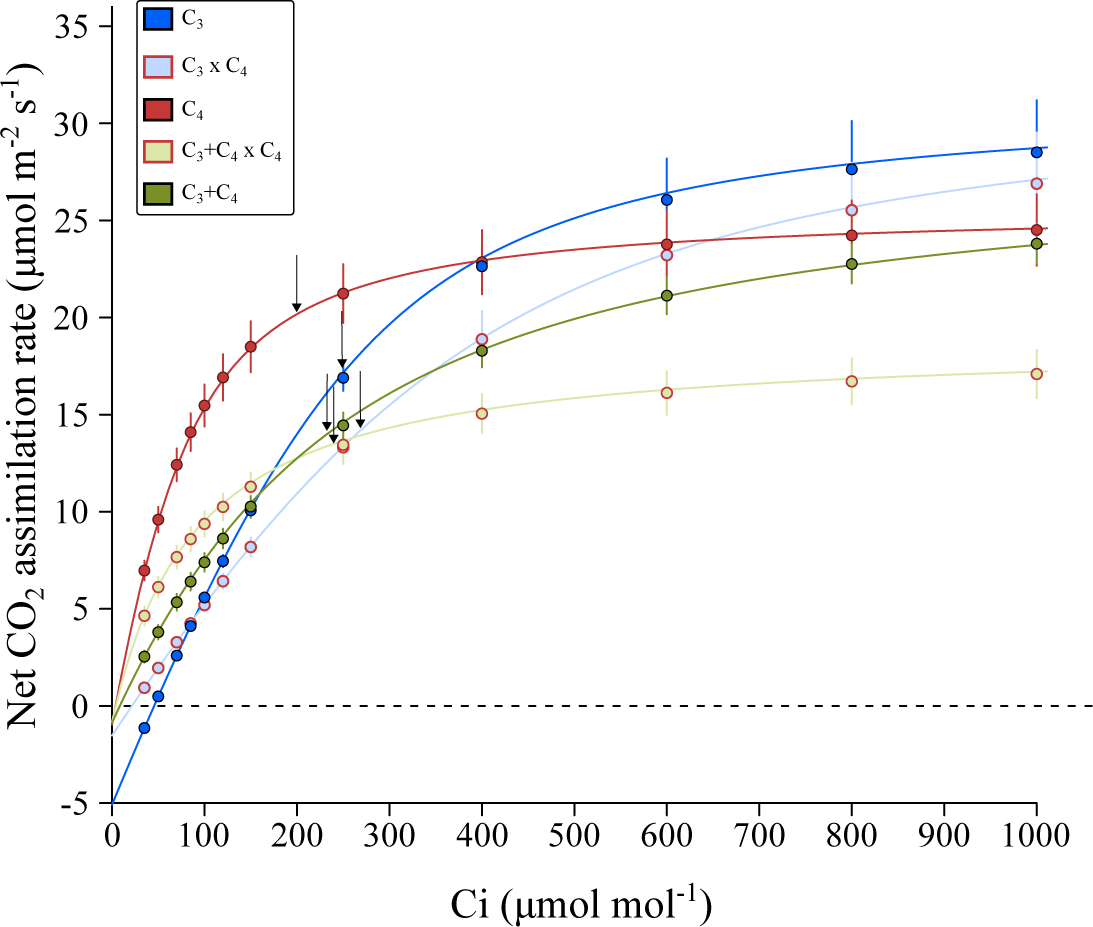
Response of net CO_2_ assimilation rate (*A*) to intercellular CO_2_ (*Ci*) of F1 hybrids and the parental photosynthetic types in *Alloteropsis semialata*. Data points are predicted values from an empirical non-rectangular hyperbola model (Bellasio et al. 2016) based on the mean parameter estimates per individual within each cross/photosynthetic type. Error bars show standard errors for the predicted values (n ≥ 3). Arrows indicate *Ci* values at reference CO_2_ = 400 µmol mol^-1^ for each cross/photosynthetic type.

**Fig. 5.**
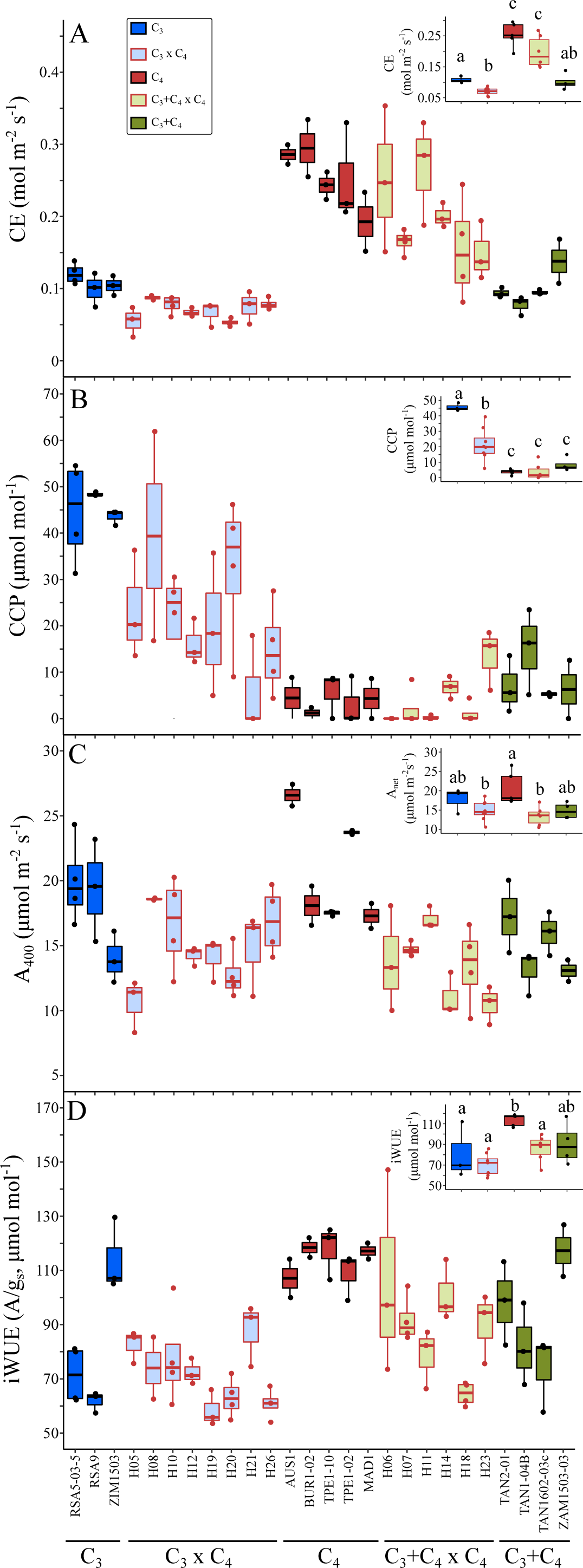
Photosynthetic performance of F1 hybrids and the parental photosynthetic types in *Alloteropsis semialata*. (A) Maximum carboxylation efficiency (CE), (B) CO_2_ compensation point (CCP), and steady-state (C) net photosynthetic rate (A_400_), and (D) intrinsic water use efficiency (iWUE, *A*/*gs*) at reference CO_2_ = 400 µmol mol^-1^. Different lower-case letters indicate statistical differences between groups (ANOVA, *p* < 0.05 post-hoc Tukey HSD; n ≥ 4).

The steady-state net CO_2_ assimilation rate at ambient CO_2_ (400 µmol mol^-1^; A_400_) was highest in two C_4_ accessions, but the differences between C_3_, C_3_+C_4_ and C_4_ plants were not significant (Fig. 5C). C_3_+C_4_ x C_4_ hybrids had the lowest A_400_, but mean values were not significantly different from C_3_ x C_4_, C_3_ and C_3_+C_4_ accessions. Stomatal conductance (*gs*) had the highest median values in C_3_ accessions, but the differences between groups were not significant (Fig. S7A). C_3_ accessions had on average 50% higher stomatal density than C_3_+C_4_ and C_4_ accessions and their hybrids (Fig. S7B). We found a positive correlation between *gs* and stomatal density (R² = 0.31, *p* < 0.01), but only after excluding the outlier C_3_ accession ZIM1503, which had the highest stomatal density, but one of the lowest *gs*. The correlation between stomatal density and vein density was only marginally significant (R² = 0.01, *p* = 0.06). The intrinsic water use efficiency (iWUE) at ambient CO_2_ was on average 38% higher in C_4_ than in other accessions, although a few C_3_, C_3_+C_4_ and C_3_+C_4_ x C_4_ individuals also showed C_4_-like values (Fig. 5D). The lowest iWUE values were observed in C_3_ and C_3_ x C_4_ individuals, but the means were not significantly different from C_3_+C_4_ and C_3_+C_4_ x C_4_ hybrids. To investigate whether changes in leaf temperature (T_leaf_) could explain the observed patterns, we inspected the variation in T_leaf_ during the A/*Ci* curves. Although T_leaf_ values up to 3°C above the median (26.3°C) were observed in a few accessions, 81% of all data points were collected within a 2°C interval (Fig. S8). There were no significant differences between groups when all points were considered, nor when only above- or below-ambient CO_2_ levels were analysed (Fig. S8). Similar patterns among cross/photosynthetic types were observed in all photosynthetic parameters after removing individuals for which the median T_leaf_ was 1°C above the median of all curves (Fig. S9).

Overall, these results show that the physiological characters are not consistently additive in the hybrids, with some individuals performing below their two parents for several traits.

## Discussion

### Partial contribution of the C_4_ cycle to carbon fixation in hybrids

In this study, we analyse the phenotypes of hybrids between individuals of the grass *Alloteropsis semialata* with different photosynthetic types. The C_3_ x C_4_ quantitatively resemble the naturally occurring C_3_+C_4_ in most anatomical and gene expression traits (Figs 2 and 3), as previously noticed (Kadereit et al. 2017). They however differ qualitatively, and minor veins and high expression of some C_4_-related genes are especially restricted to C_3_ x C_4_ individuals (Fig. 3; Table S6). These findings therefore provide additional support for the hypothesis that naturally occurring C_3_+C_4_ in *A. semialata* do not result from recent crosses between photosynthetic types (Lundgren et al. 2015, 2016), although genome analyses support a role for introgression in their distant history (Bianconi et al. 2020).

Across all crosses, we show that hybrids between C_4_ and non-C_4_ individuals of the grass *A. semialata* do not exhibit a full C_4_ physiology, as indicated by their δ^13^C, which are clearly outside the range of C_4_ values in the species (Fig. 1). This indicates that a significant fraction of CO_2_ assimilation in these hybrids occurs directly via Rubisco in the mesophyll, which implies that the genetic contribution of the C_4_ parent was insufficient to 1) completely suppress Rubisco expression and/or activity in the mesophyll cells, and/or 2) create an effective C_4_ cycle. However, the CCP of the C_3_ x C_4_ hybrids is substantially reduced compared to the C_4_ parents (Fig. 5B). While this change could theoretically result from an increase of the photorespiratory shuttle observed in many C_3_-C_4_ plants (called ‘C2’ plants; Keerberg et al. 2014; Khoshravesh et al. 2016), the expression of photorespiratory is rather unchanged or decreased in our hybrids compared to the C_4_ parents. Changes of CCP are therefore more likely linked to an increase of the C_4_ activity, which is further supported by significantly higher δ^13^C values compared to their non-C_4_ parents (Fig. 1). Our results therefore suggest that, in most hybrids, a fraction of the CO_2_ that enters the leaf is fixed by PEPC in the mesophyll and follows the C_4_ route to be refixed by Rubisco in the inner bundle sheath. Furthermore, given the C_4_-like CCP and CE values, and δ^13^C above -20‰ (in most cases), the degree of C_4_ activity is higher in C_3_+C_4_ x C_4_ than in C_3_ x C_4_ hybrids. Our findings therefore recapitulate what has been shown for interspecific hybrids between photosynthetic types in several eudicot and monocot genera (Björkman et al. 1969; Brown et al. 1985; Araus et al. 1990; Byrd et al. 1992; Brown and Bouton 1993; Oakley et al. 2014; Sultmanis 2018). The causes for this only partial contribution of the C_4_ pathway to the overall carbon assimilation in the hybrids are potentially the same in all these cases, and are probably related to the insufficient expression of components of C_4_ metabolism (Brown and Bouton 1993).

Two interrelated leaf anatomical traits were significantly different from the C_4_ individuals in both C_3_ x C_4_ and C_3_+C_4_ x C_4_ hybrids, namely the proportion of inner sheath to mesophyll tissue (IS/M) and the distance between consecutive bundle sheaths (BSD). These traits are mostly determined by the presence of minor veins in C_4_ leaves, which has been previously shown to be the major anatomical difference between C_4_ and non-C_4_ *A. semialata* (Lundgren et al. 2019). Here, significant changes towards the C_4_ phenotype in IS/M and BSD were observed in the hybrids, but realized values were still far from those observed in C_4_ accessions (Fig. 2C,D). This is particularly clear for IS/M, where values were below the mean of both parents in the two hybrid types. While minor veins are observed in the hybrids, their number per segment remains below that observed in C_4_ accessions (Table S6). This indicates that the proliferation of minor veins is a continuous trait, which can sustain a C_4_ cycle only over a threshold in this species. Since CO_2_ refixation by Rubisco takes place in the IS cells in C_4_ *A. semialata* (Ellis 1974b; Ueno and Sentoku 2006), the leaf anatomy of the hybrids might prevent an optimal coupling between carbon assimilation and reduction reactions, therefore decreasing the efficiency of the C_4_ pathway. The distribution of organelles among cell types was not quantified here, but a gene previously linked to the relocation of organelles to BS cells (Wang et al. 2017) is more highly expressed in C_4_ accessions and upregulated in some hybrids compared to their non-C_4_ parents, which might suggest that mitochondria are partially relocated to their BS. The size of the OS cells might also play a role in creating an effective C_4_ cycle, as suggested by the large differences between C_3_ and C_4_ accessions (Fig. 2E). Few other C_4_ species have an extra layer of BS cells outside of the BS cells containing chloroplasts, and the presence of an OS may represent an increased resistance to the C_4_ acid shuttle (Lundgren et al. 2014). In the grass genus *Neurachne* that presents a similar C_4_ anatomy, the smallest OS are also observed in C_4_ individuals, with intermediate species presenting intermediate OS cell sizes (Khoshravesh et al. 2020), supporting the idea that OS cells must be reduced during C_4_ evolution. Here, the OS cells were significantly reduced in the hybrids relative to their non-C_4_ parents, with most C_3_+C_4_ x C_4_ individuals and a few C_3_ x C_4_ having OS cell sizes within the C_4_ range (Fig. 2F), which suggests that this character might have played a minor role, if any, in reducing the efficiency of the C_4_ acid shuttle in the hybrids.

C_4_ activity could also have been limited by gene expression and protein activity. Despite significant increases in the transcript abundance of core C_4_ genes, particularly the genes encoding PEPC (Fig 4), it is possible that the insufficient expression levels of some genes might have reduced the efficiency of the C_4_ cycle, for example by decreasing the rate of carboxylation and/or decarboxylation reactions. Furthermore, the lack of proper compartmentation of the expression/activity of these genes due to a non-tissue-specific expression of C_3_ or C_3_+C_4_ alleles could have impaired the formation of intermediate metabolite pools, therefore preventing the operation of an efficient C_4_ cycle (Brown and Bouton 1993; Ermakova et al. 2020). Finally, the lack or insufficiency of post-transcriptional and post-translational modifications might also have affected the realized enzyme activity. This might be particularly relevant in the case of hybrids, where the cell environment may be substantially modified by pleiotropic effects emerging from the expression of two divergent genomes. Detailed analyses of cellular localization and activity of C_4_ enzymes coupled to CO_2_ labelling studies are necessary to identify the major determinants of the phenotype observed in the hybrids reported here.

### Photosynthetic performance of plants expressing partial C_4_ traits and implications for the model of C_4_ evolution

The current model of C_4_ evolution assumes that the sequential acquisition of certain features, such as Rubisco activity in bundle sheath cells and upregulation of core C_4_ enzymes, increases photosynthetic output and consequently fitness in an environment that favours the C_4_ physiology (Monson and Moore 1989; Sage 2004; Heckmann et al. 2013; Malmmann et al. 2014). However, this Mt. Fuji-like fitness landscape (Heckmann et al. 2013) is conditional on the presence of anatomical enablers; when these are lacking, the C_4_ phenotype is not accessible because intermediate states have lower fitness than the C_3_ state (Heckmann 2016). The two groups of intraspecific hybrids generated in this study provide an opportunity to examine this prediction while controlling for phylogenetic effects, with C_3_ x C_4_ and C_3_+C_4_ x C_4_ plants being proxies for the upregulation of C_4_ traits at an early and intermediate stage, respectively, along C_4_ evolution. Here, we show that the carboxylation efficiency of C_3_ x C_4_ hybrids is lower than that of both parents (Figs 5, 6A), despite significant changes towards the C_4_ phenotype in leaf anatomy and gene expression. In C_3_+C_4_ x C_4_ hybrids, however, carboxylation efficiency was increased in comparison to the C_3_+C_4_ parent, and was not significantly different from C_4_ accessions. Such disparate effects of upregulating C_4_ components in a non-C_4_ background are compatible with the initial prediction that fitness gains are conditional on the presence of enabling factors, which here might have been present in C_3_+C_4_ parents, but lacking in the C_3_ plants. Such changes would have acted as a switch that permitted a full C_4_ pathway to operate, and therefore increase photosynthetic performance.

Note however that such gains in photosynthetic performance are restricted to conditions of low CO_2_ concentration in the leaf (Fig. 4). In fact, at ambient CO_2_ levels, several individuals of both hybrid groups had lower photosynthetic rates than both parents, suggesting that the partial upregulation of C_4_ components, all at once, might have negative consequences, possibly due to pleiotropic effects (e.g. C_4_ enzymes competing for reducing power and ATP with other cell reactions). This in turn supports the idea that the changes towards a full C_4_ physiology must be built sequentially upon preexisting characteristics that enable the next change to provide a fitness advantage (Heckmann 2016). The fertility of the F1 hybrids reported here is not known yet, although evidence of rare gene flow between photosynthetic types in the wild suggest that backcrossing is possible (Olofsson et al. 2016, 2021; Bianconi et al. 2020). If a F2 population can be produced, characterization of the photosynthetic performance of such individuals expressing C_4_ components independently, particularly under conditions expected to favour C_4_ plants, including long-term exposure to low CO_2_ levels, should provide further insights on whether C_4_ evolution must follow a particular sequence of events for it to be viable.

## Conclusions

In this study, we characterize the phenotype of hybrids between different photosynthetic types in the grass *Alloteropsis semialata* to investigate how the upregulation of components of the C_4_ trait affects photosynthetic performance. We show that the hybrids have in most cases anatomical traits and gene expression patterns that are intermediate between those of the parents, and this leads to C_4_ activity that is equally intermediate between the two parents. The physiological benefits of a partial C_4_ metabolism in the hybrids appear only at low CO_2_ levels; at ambient CO_2_, there is no evidence of enhanced photosynthetic performance in the hybrids relative to their non-C_4_ parents. Some hybrid individuals in fact perform worse than both parents at ambient CO_2_, and this possible hybrid depression could explain the lack of a clear hybrid zone in regions where the distributions of C_4_ and non-C_4_ *A. semialata* lineages overlap. Overall, our results support the hypothesis that photosynthetic gains arising from the upregulation of C_4_ features are conditional on coordinated changes in leaf anatomy and biochemistry. Future studies with the F2 offspring, where the different C_4_ components can be segregated, will be able to pinpoint the key genetic and phenotypic changes that lead to fitness gains, providing a unique opportunity to experimentally test long-standing hypotheses about the evolution of C_4_ photosynthesis.

## Author contributions

MEB, CPO and PAC designed the study. MEB and LTB phenotyped the plants. EVC, GS and LTD PCR-genotyped the plants. GS and VM generated transcriptome data. ES produced the hybrids. MEB analysed the data. MEB and PAC wrote the manuscript with the help of all co-authors.

## Acknowledgements

This work was funded by the European Research Council (grant ERC-2014-STG-638333) and the Royal Society (grant RGF\EA\181050). M.E.B is funded by a Royal Society grant (RGF\EA\ 181050), and P.A.C. by a Royal Society University Research Fellowship (grant URF\R\180022).

## Data availability

Newly generated RNA sequencing datasets were deposited in the NCBI SRA database under Bioproject PRJNA752516. Scripts used for the genotyping analysis are available at https://github.com/matheusbianconi/RNAgenotyping_hybrids

## Supplementary data

**Fig. S1.**
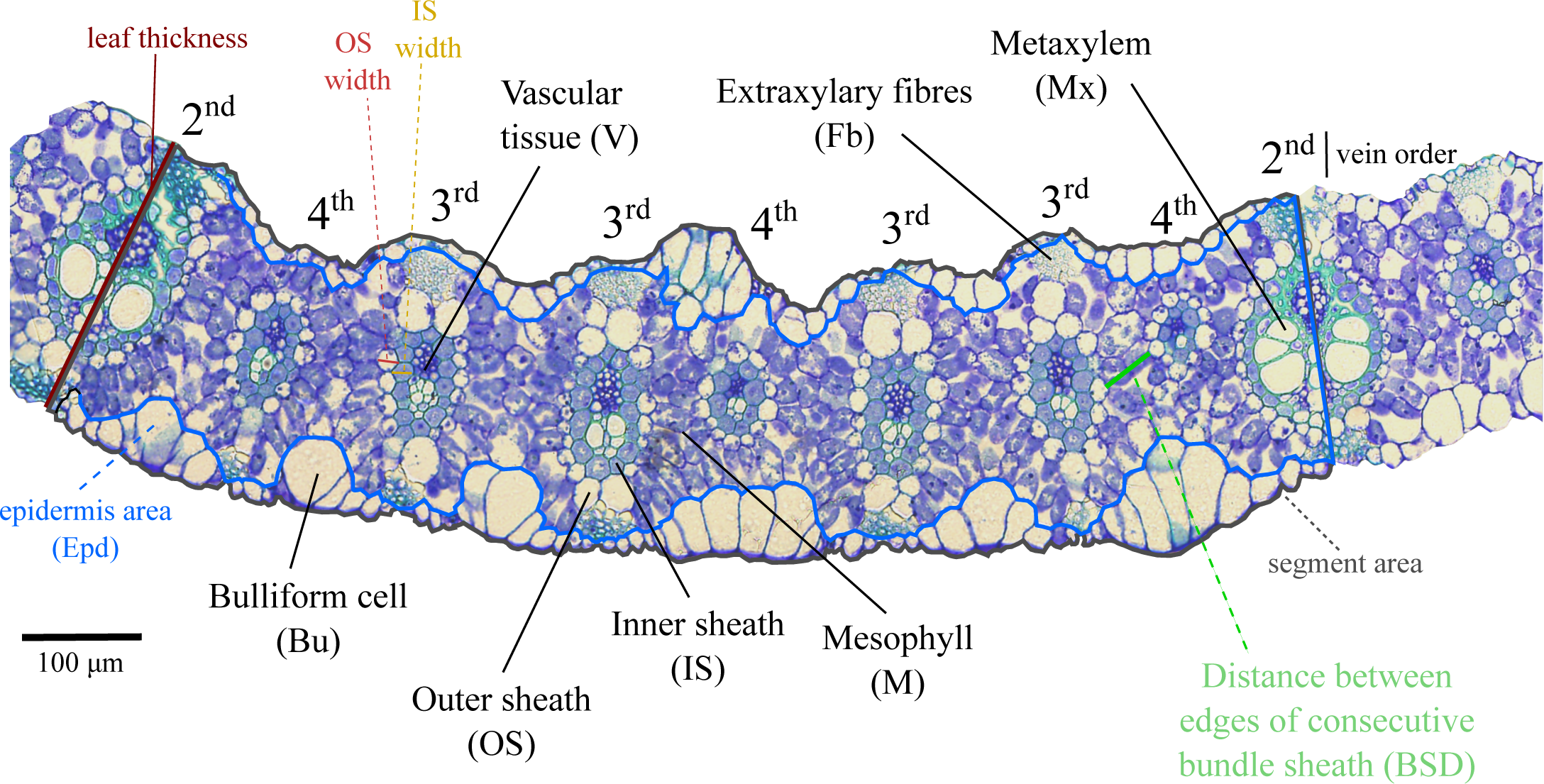
Leaf anatomical variables measured in this study.

**Fig. S2.**
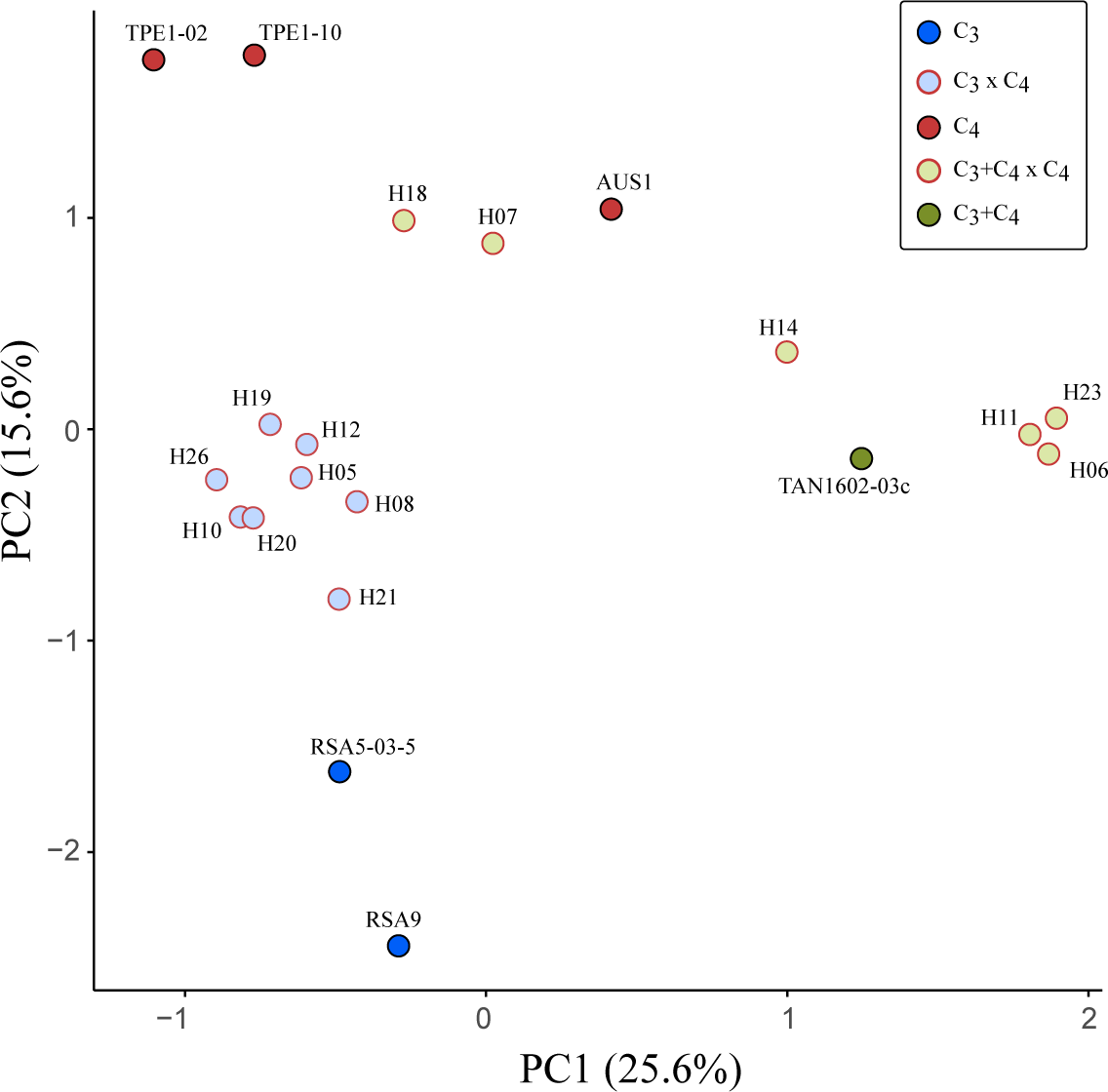
Principal component analysis on 11,988 genes from the chromosome-level genome assembly of *Alloteropsis semialata*.

**Fig. S3.**
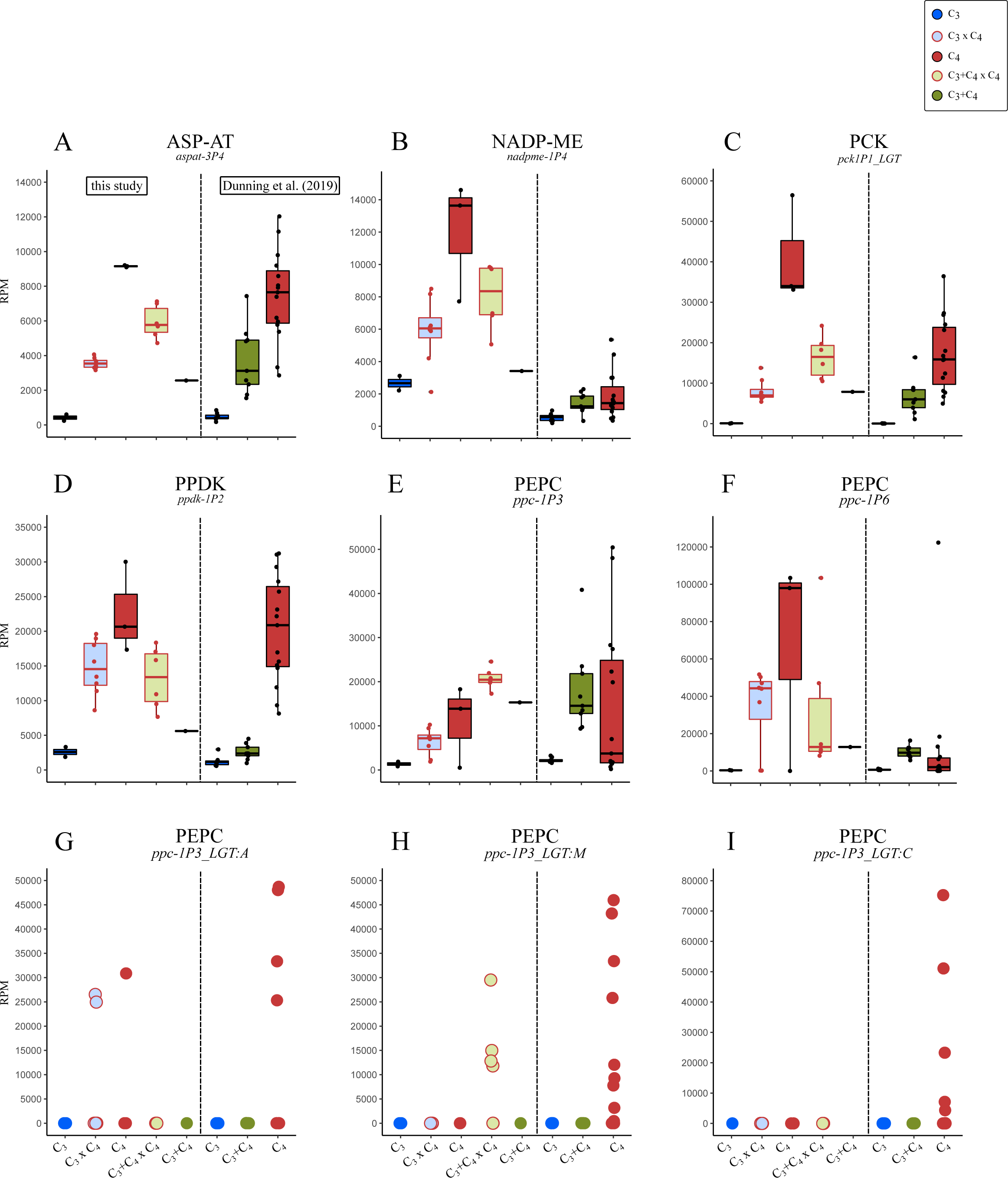
Transcript abundance in reads per million (RPM) of selected genes encoding core C_4_ enzymes in *Alloteropsis semialata*: (A) aspartate aminotransferase (ASP-AT, gene *aspat-3P4*), (B) NADP-dependent malic enzyme (NADP-ME, gene *nadpme-1P4*), (C) phosphoenolpyruvate carboxykinase (PCK, gene *pck1P1_LGT*), (D) pyruvate phosphate dikinase (PPDK, gene *ppdk-1P2*), and (E-I) phosphoenolpyruvate carboxylase (PEPC) genes. Transcript abundance computed for *A. semialata* samples extracted from Dunning et al. (2019a) are shown on the right for each gene.

**Fig. S4.**
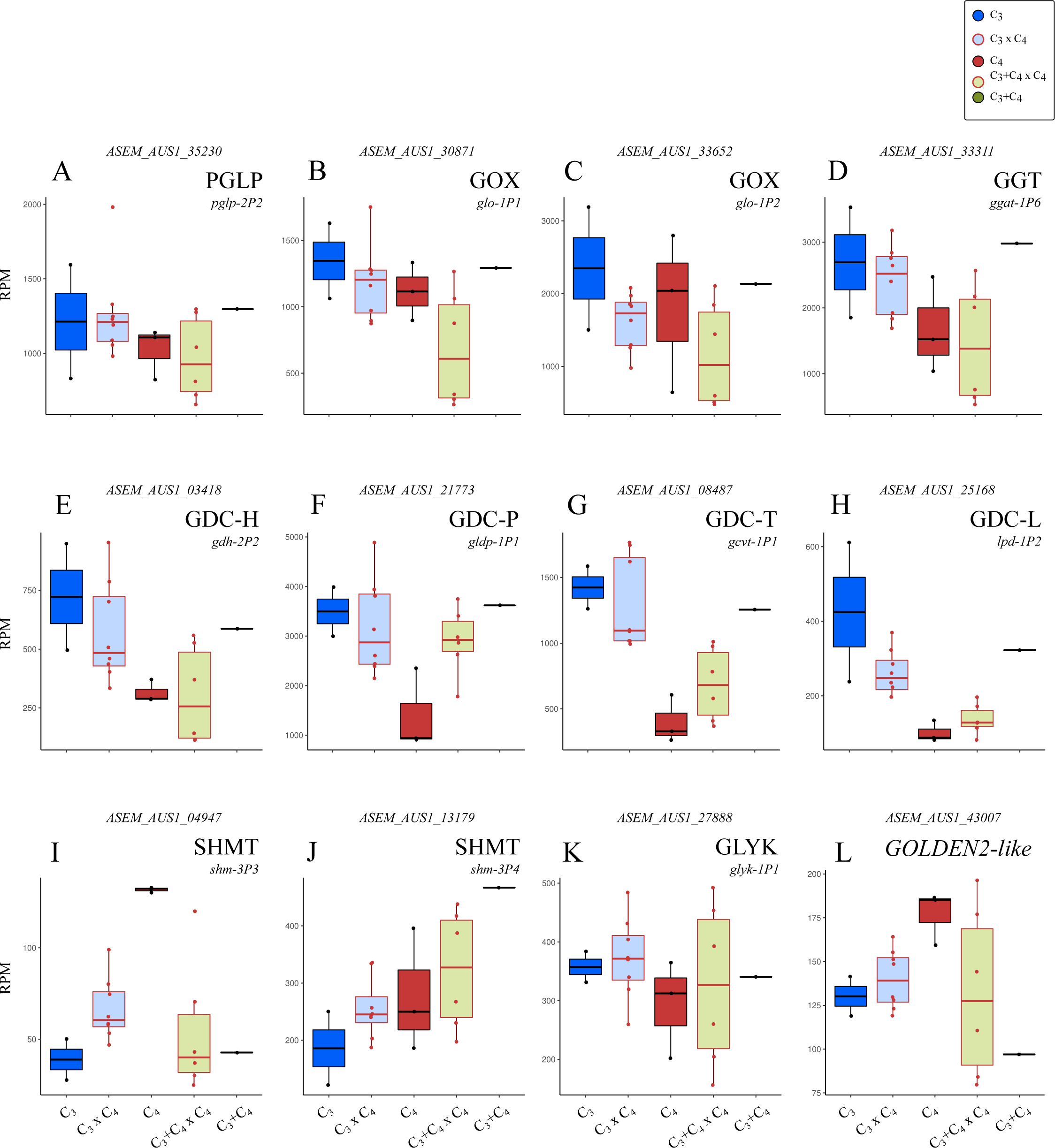
Transcript abundance of selected gene families with a role in photorespiration (A-K), and the gene encoding the transcription factor GOLDEN2-like (L) in F1 hybrids and the parental photosynthetic types in *Alloteropsis semialata*. (A) 2-phosphoglycolate (2-PG) phosphatase (PGLP), (B-C) flavin mononucleotide (FMN)-dependent glycolate oxidase (GOX), (D) glutamate:glyoxylate aminotransferase (GGT), (E-H) glycine decarboxylase (GDC) complex proteins -H, -P, -T and -L, (I-J) serine hydroxymethyltransferase (SHMT), (K) glycerate 3-kinase (GLYK). Transcript abundance in reads per million mapped reads (RPM).

**Fig. S5.**
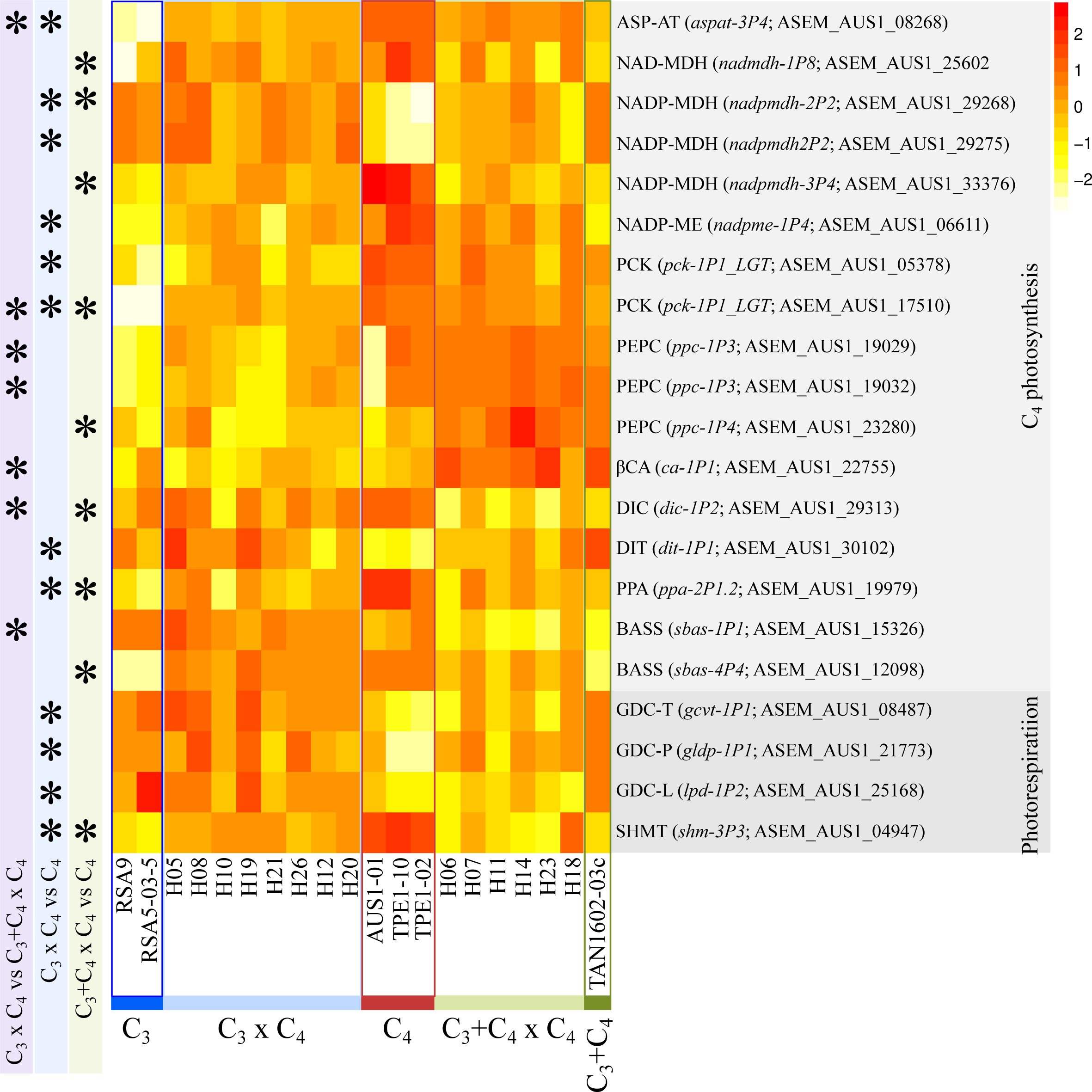
Heat map of differentially expressed genes related to C_4_ photosynthesis and photorespiration. Asterisks indicate significant DE genes (*p* < 0.05) for comparisons between hybrid types (C_3_ x C_4_ vs C_3_+C_4_ x C_4_) and between each hybrid type and the C_4_ type. Only DE genes with at least two-fold change and base count > 500 are shown. Count data was transformed using the VST function of DESeq2 and scaled by row. See table S9 for full gene annotation

**Fig. S6.**
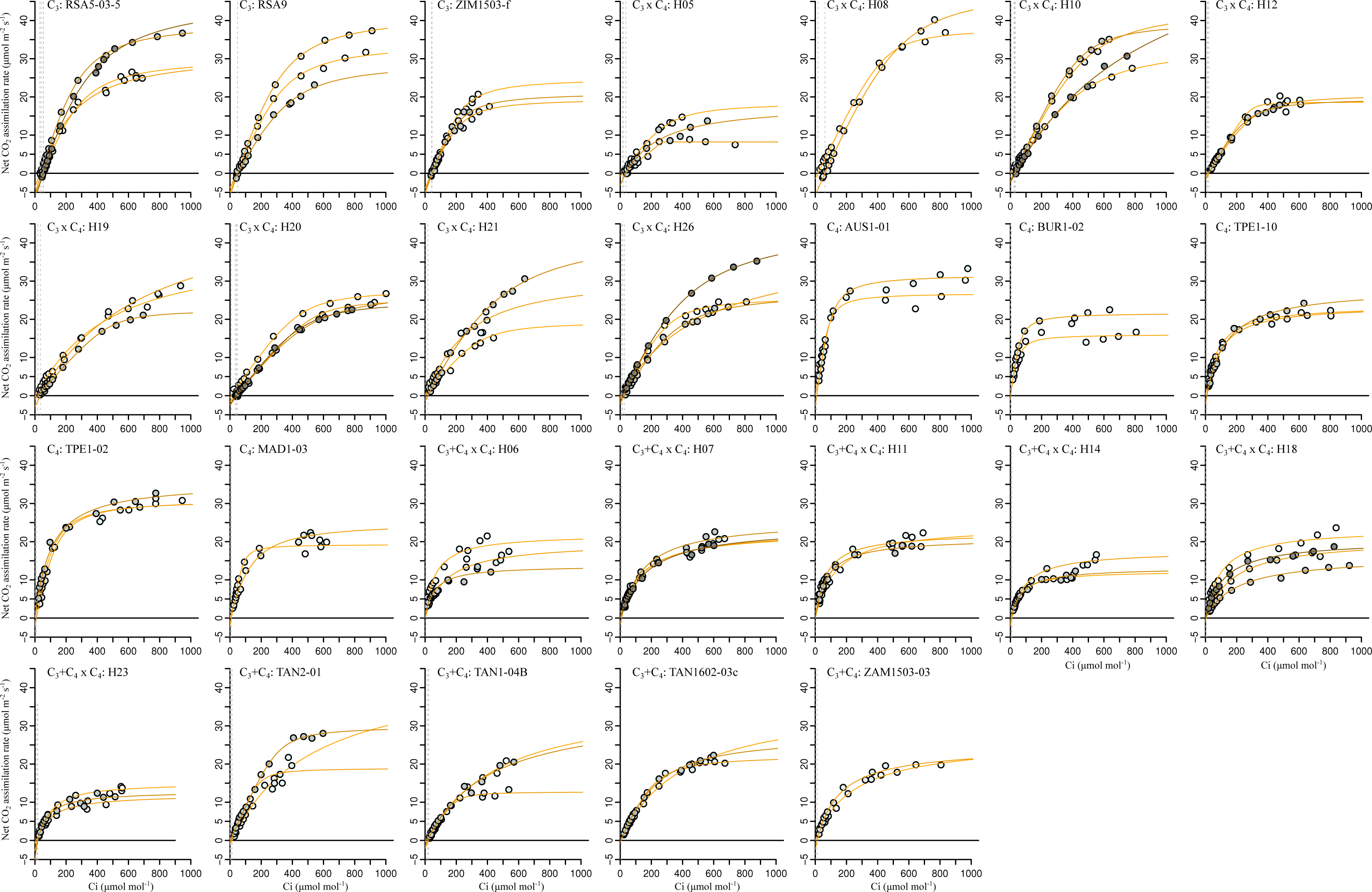
Photosynthetic response to intercellular CO_2_ (*A*/*Ci*) of F1 hybrids and the parental photosynthetic types in *Alloteropsis semialata*. Panels contain all *A*/*Ci* curves collected for each accession, with individual curves coloured with different shades of grey. Accession name and cross/photosynthetic type are indicated on the top left. Vertical dashed lines indicate the CO_2_ compensation point of each curve.

**Fig. S7.**
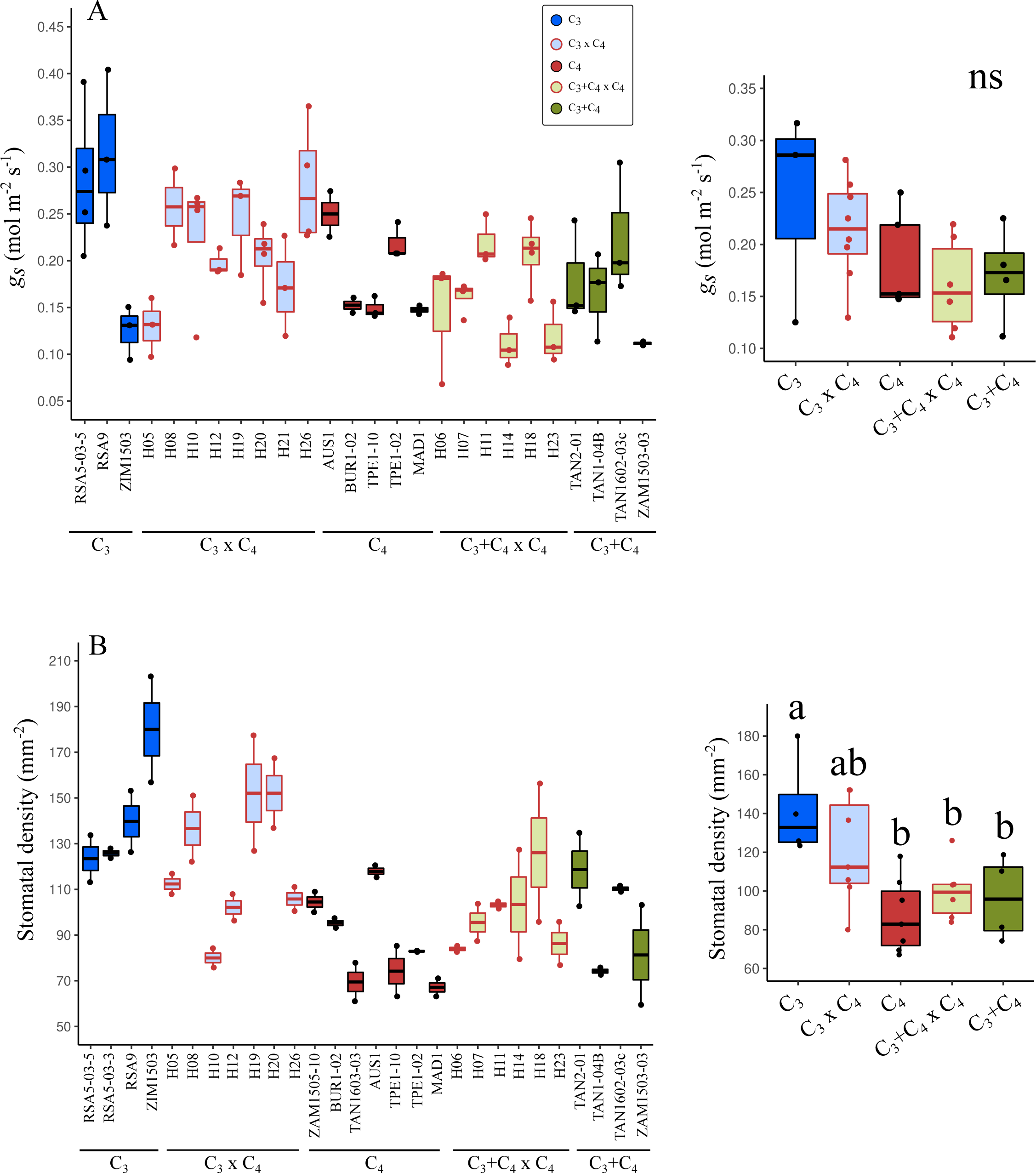
Leaf stomata variables. (A) Steady-state stomatal conductance (*gs*) at 400 μmol mol^-1^ (n = 2-4 leaves per accession). (B) Stomatal density on the abaxial side of the leaves (n = 2 leaves per accession, with stomata counts averaged from 5 fields per leaf; field area = 0.38 mm^2^). Data points on the right are the means per accession within each cross/photosynthetic type, and different lower-case letters indicate statistical differences between groups (ANOVA, *p* < 0.05 post-hoc Tukey HSD; n ≥ 3).

**Fig. S8.**
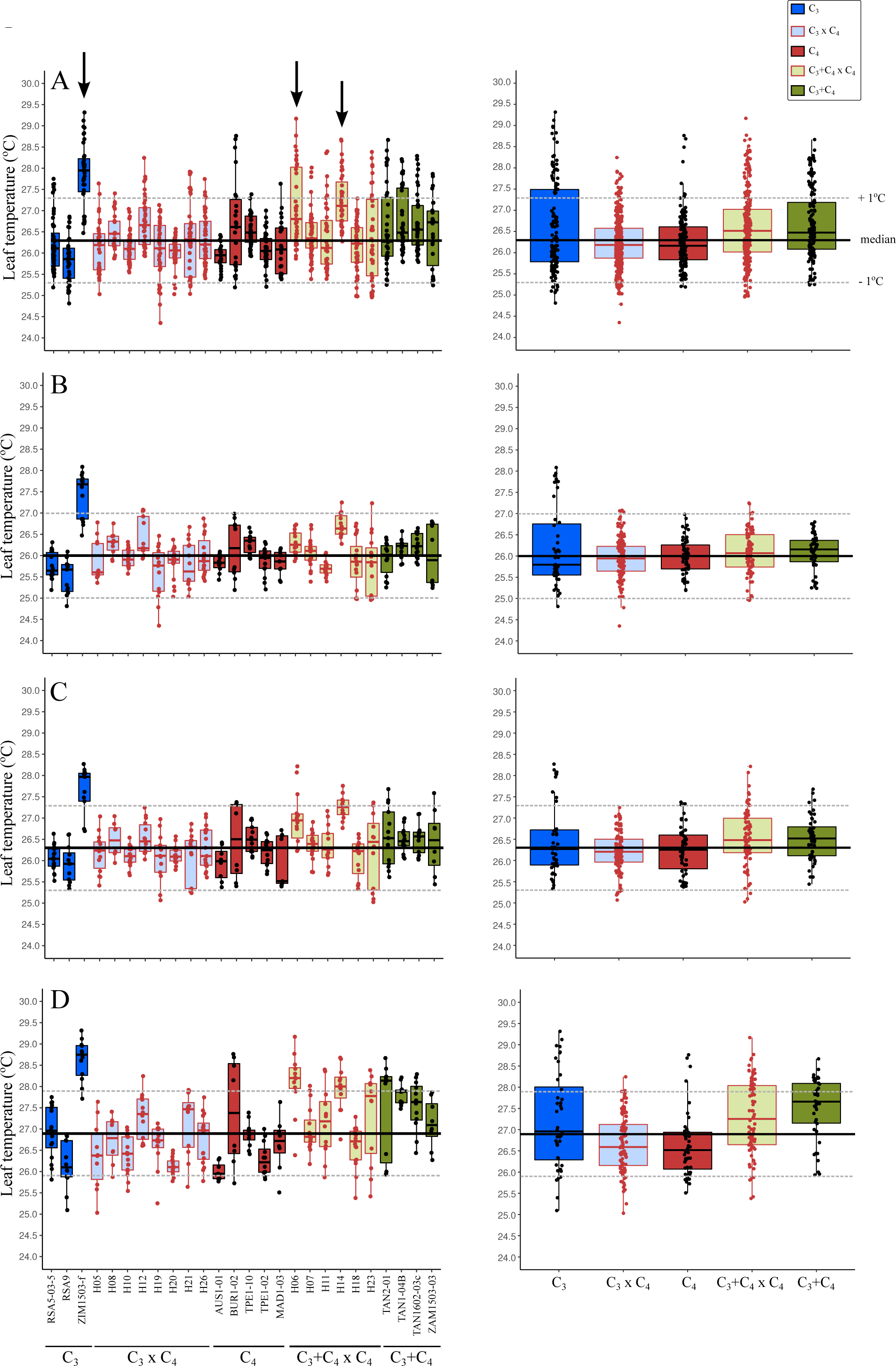
Leaf temperature variation during *A*/*Ci* curves. Data points are individual *A*/*Ci* measurements for each plant (left panels) or with individuals grouped into cross/photosynthetic types (right panels). (A) All measurements, and measurements collected at reference CO_2_ (B) < 100 µmol mol^-1^, (C) between 100 and 400 µmol mol^-1^, and (D) > 400 µmol mol^-1^. Arrows indicate outlier individuals that were removed for the analysis in Fig. S9.

**Fig. S9.**
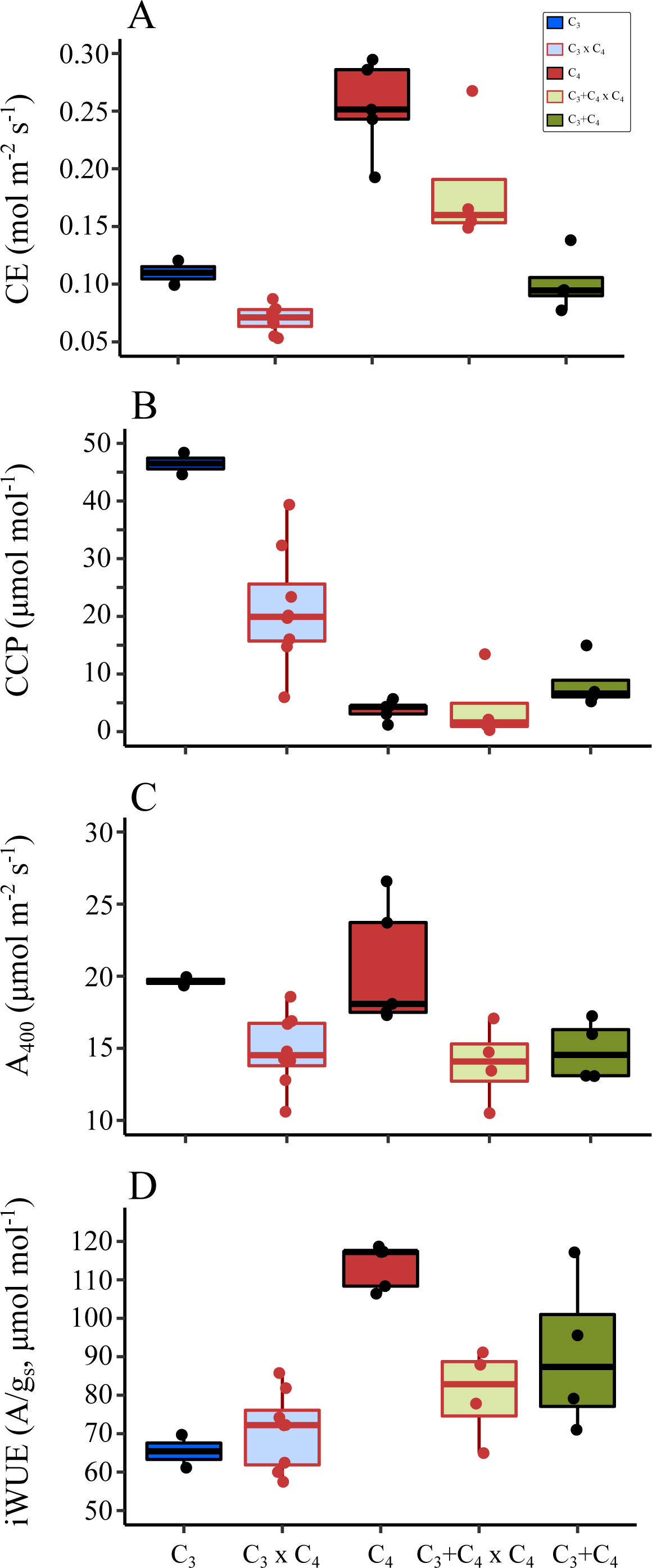
Photosynthetic performance of F1 hybrids and the parental photosynthetic types in *Alloteropsis semialata* after removing outlier accessions (i.e. with T_leaf_ 1°C above the median). (A) Maximum carboxylation efficiency (CE), (B) CO_2_ compensation point (CCP), and steady-state (C) net photosynthetic rate (A_400_), and (D) intrinsic water use efficiency (iWUE, *A*/*gs*) at reference CO_2_ = 400 µmol mol^-1^.

---

**Table S1.** Sample information.

**Table S2.** Genotyping using PCR/Sanger-sequencing.

**Table S3.** List of primer sequences used for genotyping.

**Table S4.** RNA-seq markers selected for genotyping analysis I (photosynthetic type).

**Table S5.** RNA-seq markers selected for genotyping analysis II (pollen parent).

**Table S6.** Raw leaf anatomy data.

**Table S7.** Normalized transcript abundance for the co-ortholog gene set.

**Table S8.** Normalized transcript abundance for the *A. semialata* complete genome gene set.

**Table S9.** Differential expression analysis between F1 hybrids and the C_4_ parental type.

**Table S10.** Gene ontology enrichment analyses on genes differentially expressed between F1 hybrids and the C_4_ parental type.

**Table S11.** Raw A/*Ci* data.

**Table S12.** Estimated photosynthetic parameters and steady-state measurements.

## Supplementary Methods

### Global transcriptome analyses

We performed differential expression (DE) analyses using the raw counts obtained after mapping the RNA-seq datasets to the coding sequences extracted from the chromosome-level genome assembly of *A. semialata* (Dunning et al. 2019; see main text; total number of genes = 45,145). To detect genes that were differentially expressed between each of the hybrid types and the parental types, we used the R package *DESeq2* (Love et al. 2014). Due to the lack of sufficient replicates for the C_3_ and C_3_+C_4_ parental types, we restricted our analyses to the comparisons between hybrid types (C_3_ x C_4_ vs C_3_+C_4_ x C_4_), and between each of these and the C_4_ type. Only genes with more than 10 counts across all accessions were retained for the DE analysis (total = 29,200 genes). For each comparison, we used a false discovery rate (FDR) of 0.05 as cut-off value. We then investigated whether each of the sets of DE genes were associated with any particular metabolic function using a gene ontology (GO) enrichment analysis with the R package *clusterProfiler* (Yu et al. 2012). We first used Orthofinder v2.5.2 (Emms and Kelly 2019) with default parameters to identify the corresponding orthologs of *A. semialata* in the closely related grass species *Setaria italica* (v2.2), *Sorghum bicolor* (v3.1.1) and *Oryza sativa* (v7.0; genomes extracted from Phytozome v13; Goodstein et al. 2012). We extracted the GO annotations for the three genomes using the Biomart tool of Phytozome 13, and transferred the annotations to the *A. semialata* gene set using the orthology information obtained from Orthofinder. We then performed a GO enrichment analysis on the DE gene set resulting from each of the three comparisons using the *enricher* function of *clusterProfiler* with a *p*-value cut-off of 0.05. The identity of C_4_- and photorespiration-related genes in the genome of *A. semialata* was extracted from the annotations used in Bianconi et al. (2018) and Dunning et al. (2019).

